# Genome-wide prediction and integrative functional characterization of Alzheimer’s disease-associated genes

**DOI:** 10.1101/2021.02.09.430536

**Authors:** Cui-Xiang Lin, Hong-Dong Li, Chao Deng, Weisheng Liu, Shannon Erhardt, Fang-Xiang Wu, Xing-Ming Zhao, Jun Wang, Daifeng Wang, Bin Hu, Jianxin Wang

## Abstract

The mechanism of Alzheimer’s disease (AD) remains elusive, partly due to the incomplete identification of risk genes. We developed an approach to predict AD-associated genes by learning the functional pattern of curated AD-associated genes from brain gene networks. We created a pipeline to evaluate disease-gene association by interrogating heterogeneous biological networks at different molecular levels. Our analysis showed that top-ranked genes were functionally related to AD. We identified gene modules associated with AD pathways, and found that top-ranked genes were correlated with both neuropathological and clinical phenotypes of AD on independent datasets. We also identified potential causal variants for genes such as *FYN* and *PRKAR1A* by integrating brain eQTL and ATAC-seq data. Lastly, we created the ALZLINK web interface, enabling users to exploit the functional relevance of predicted genes to AD. The predictions and pipeline could become a valuable resource to advance the identification of therapeutic targets for AD.

## Introduction

Alzheimer’s disease (AD) is a complex and progressive neurodegenerative disorder that accounts for the majority of all dementia cases^1^. Its clinical symptoms include progressive memory loss, personality change, and impairments in thinking, judgment, language, problem-solving, and movement^2^. The two neuropathological hallmarks of AD are extracellular amyloid-β (Aβ) plaques and intracellular neurofibrillary tangles (NFTs), which are known to contribute to the degradation and death of neurons in the brain^3^. The number of patients with AD worldwide is currently rising. Specifically, it is estimated that approximately 50 million people are currently living with AD or other forms of dementia, and this number is expected to increase to over 152 million by 2050^1^. AD not only causes suffering in both patients and their families but also places a severe burden on society. However, the drug development for AD is slowly progressing^4^, partly due to the incomplete understanding of the neuropathological mechanisms.

AD is partly caused by genetic mutations^4^. Its two subtypes, *i*.*e*., early-onset AD (EOAD, onset age before 65 years) and late-onset AD (LOAD, onset age later than 65 years), have different genetic risk factors. In EOAD, rare mutations in *APP, PSEN1* and *PSEN2* have been identified^4^. LOAD is markedly more complex, with *APOE* being a well-known risk gene for this subtype. Most known or putative AD-associated genes were discovered through genome-wide association studies (GWAS). Previously, GWAS identified *CLU, CR1*, and *PICALM*, along with approximately 20 more genes^4^. In addition, network approaches are used to identify AD-associated molecular networks or pathways. For example, a module-trait network approach was proposed and applied to identify gene coexpression modules that were associated with cognitive decline^5^, while a large-scale proteomic analysis identified an energy metabolism-linked protein module, strongly associated with AD pathology^6^. However, a large proportion of the phenotypic variances in AD cannot be explained by known risk genes^7, 8, 9^, which suggests additional AD-associated genes that remain to be discovered. Since experimental approaches are often time consuming and expensive, computational approaches provide a promising alternative to discovering AD-associated genes.

Previous studies have shown that functional gene networks (FGNs) are promising for predicting disease-associated genes^10, 11^. In a FGN, a node represents a gene and the edge between two genes represents the co-functional probability (CFP) that the two genes take participate in the same biological process or pathway^12^. For example, Guan *et al*. constructed a global (*i*.*e*., non-tissue specific) FGN for mice, and identified *Timp2* and *Abcg8* as two novel genes associated with bone-mineral density^13, 14^. Using the same network, Recla *et al*. discovered *Hydin* as a new thermal pain gene^13, 14^. Because gene interactions might be rewired in different tissues, global networks cannot reveal the differences of gene networks among tissues. To address this limitation, tissue-specific networks have been proposed to more accurately capture gene interactions in tissues. Greene *et al*. established 144 human tissue-specific networks and investigated these networks for the interpretation of gene functions and diseases^15^. Using the brain-specific network^15^, Krishnan *et al*. predicted disease genes for autism spectrum disorder^11^. By leveraging the functional genomic data of model species with similar genetic backgrounds, including mice and rats, a human brain-specific network was constructed, and its application to the identification of brain disorder-associated genes was illustrated in our previous work^16^.

Because AD is a brain disorder with genetic contributions, we hypothesized that brain-specific FGNs are informative for predicting AD-associated genes. It should be pointed out that our predictions of AD-associated genes do not indicate any causality, that is, the predicted genes may be either directly or indirectly associated with AD. To build models for AD-associated gene prediction, we first compiled AD-associated genes from multiple resources. These genes were used as positives for training models. We proposed a functional enrichment-based approach to identify negative genes that are not likely associated with AD. Next, we obtained ten brain-specific FGNs from the GIANT^15^ and BaiHui^16^ databases. After assessing the predictivity of each network by cross-validation of state-of-the-art machine learning models, we built a final model for predicting AD-associated genes through an optimal selection of networks and machine learning methods. We scored all the other human genes that were not used in model training for their association with AD. We created a pipeline to evaluate top-ranked novel candidate genes by interrogating multiple biological networks. We then identified gene modules from an AD-related network. We assessed the association of these modules and top-ranked genes with AD-related phenotypes, including Consortium to Establish a Registry for Alzheimer’s Disease (CERAD) score, Braak stage, and clinical dementia rating (CDR) on an independent dataset. We next identified a set of genes by combining our predictions and seven types of genomic evidence. We further identified potential variants that may affect the expression of prioritized genes. Lastly, we developed the ALZLINK web interface to enable the expoitation of predicted AD-associated genes. The resulting predictions and pipeline could be valuable to advance the identification of risk genes for AD.

## Results

### Prediction of AD-associated genes

Our approach leverages machine learning and a brain FGN to predict AD-associated genes. The approach consists of three main components: compilation of AD-associated (positive) and non-AD (negative) genes, construction of a feature matrix based on a brain FGN, and prediction of AD-associated genes using machine learning models (Fig. 1). We first compiled a set of AD-associated genes and non-AD genes to train models (see the *Methods* section; Supplementary Note 1). We showed that the negative genes were superior to those selected by the random sampling approach (Supplementary Fig. 1) and that the negative genes were poorly associated with AD (Supplementary Fig. 2). In addition, we tested their enrichment in three AD-related gene sets associated with cognitive decline (the m109 module with 390 genes)^5^, amyloid-beta (15 genes), and Tau pathology (28 genes)^17^ respectively, from two recent studies^5, 17^. The results showed that the negative genes were not enriched in any of the three modules or pathways (p-values = 0.99, 1, 1 respectively). Next, we extracted a feature matrix for the positive and negative genes based on FGNs. For each gene (positives, negatives, or the other genes), its CFPs with the positive genes in the network were collected into a 147-dimensional feature vector. We considered the 10 collected brain FGNs (nine from GIANT and one from BaiHui) and evaluated their ability to predict AD-associated genes using state-of-the-art machine learning methods, including LR, SVM, RF, and ExtraTrees, which were shown to be promising in a previous study^18^. We found that the network in the BaiHui database achieved the best performance based on the four methods tested and that ExtraTrees performed better than the other methods in terms of both the area under the receiver operating characteristic curve (AUROC) and the area under the precision-recall curve (AUPRC) (Fig. 2A; Supplementary Fig. 3-5). Finally, we selected this network in combination with ExtraTrees to construct the model for predicting AD-associated genes.

**Fig. 1.**
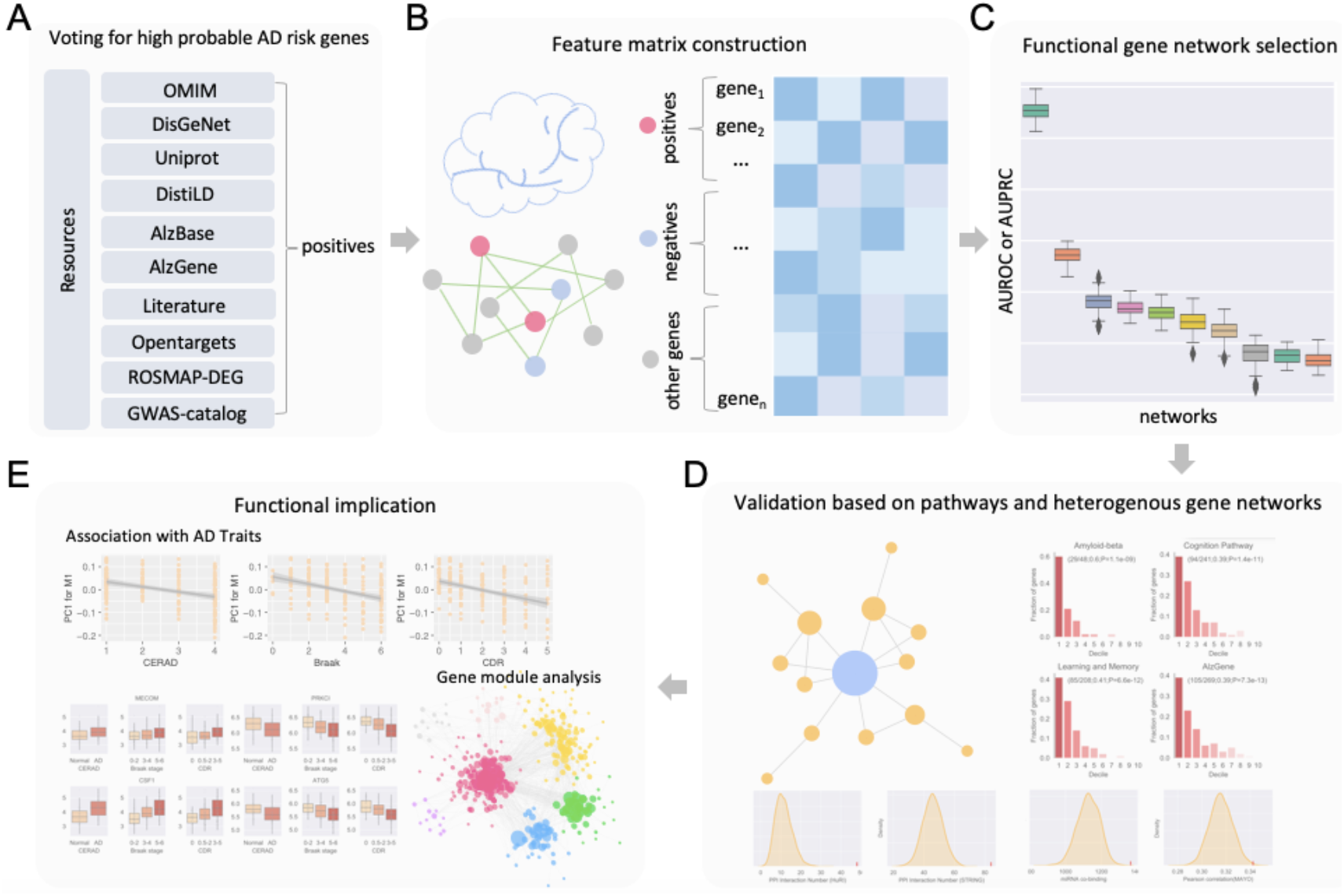
Overview of the method for genome-wide prediction of AD-associated genes and their functional characterization. **A** Selection of AD-associated genes. 147 AD-associated genes were compiled from various resources, including AD-associated genes from OMIM, DisGeNet, Uniprot, DistiLD, AlzBase, AlzBase, AlzGene, literature, Open Targets, ROSMAP-DEG and GWAS-catalog. The gene that was present in at least two resources was selected. The AD-associated genes as well as potential positive genes inferred with a functional enrichment method were then removed from the full set of all human genes. The remaining genes were treated as non-AD genes (negatives). **B** Brain specific functional gene networks (FGNs) were used for feature matrix construction. For each gene, its cofunction probabilities with the 147 positive genes in the network were extracted as features. Thus, each gene was characterized by a 147-dimensional vector. **C** Selection of brain FGNs. We compared the ten networks collected for their predictivity of AD-associated genes with machine learning approaches. An optimal network was selected. **D** Validation. Predicted AD-associated genes were validated by AD-related pathways and various gene networks, including coexpression networks, protein-protein interaction networks, miRNA-target binding networks, transcriptional regulatory networks. **E** Functional implication in AD. The associations of the top predicted genes with AD-related phenotypes were evaluated. Gene modules from an AD-related network were identified.

**Fig. 2.**
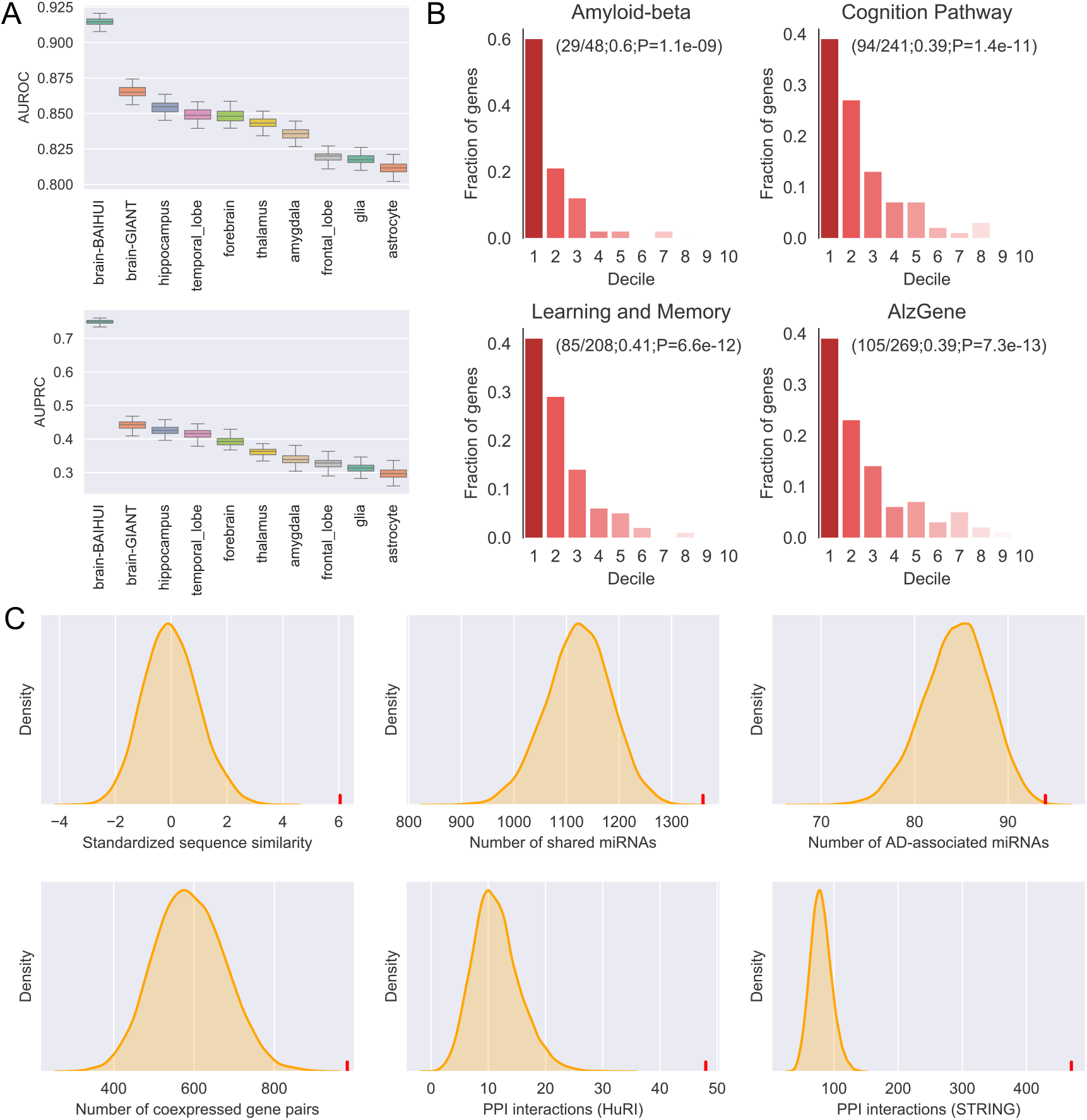
Model performance and statistical evaluation based on AD-related pathways and various gene networks. **A** Comparison of ExtraTrees models built from different functional gene networks in terms of AUROC and AUPRC based on cross-validation. **B** Enrichment of the genes ranked in the first decile in the four AD-associated gene sets or pathways with the decile enrichment test (described in Methods). **C** Validation of the top-ranked genes based on their sequence similarity, the number of shared miRNAs, the number of AD-associated miRNAs they can bind to, the number of coexpressed gene pairs, the number of interactions with AD-associated genes in HuRI and STRING. In all the subplots, the red vertical line and the distribution in yellow indicate the results for our top-ranked genes and randomly selected genes, respectively.

We performed five-fold cross-validation with ExtraTrees. Each of the five models established during cross-validation was used to score all other human genes that were not included in the training dataset. To achieve robust predictions, we repeated the cross-validation 100 times and calculated an average score for each gene. The average AUROC and AUPRC based on cross validation are 0.91 and 0.76, respectively, suggesting the model is accurate. A higher score indicates that a gene is more likely to be associated with AD. The scores for predicted genes are provided in our developed web interface (www.alzlink.com). Our literature search showed that 12 of the top-ranked 20 genes were likely associated with AD with some evidence (Supplementary Table 1), suggesting that our model has captured molecular signature of AD and makes confident predictions. Note that our prediction for AD-associated genes was based on only the machine learning model; the subsequent analysis such as enrichment, coexpression, and PPI relatedness was used separately to evaluate the association of predicted genes with AD.

**Table 1.**
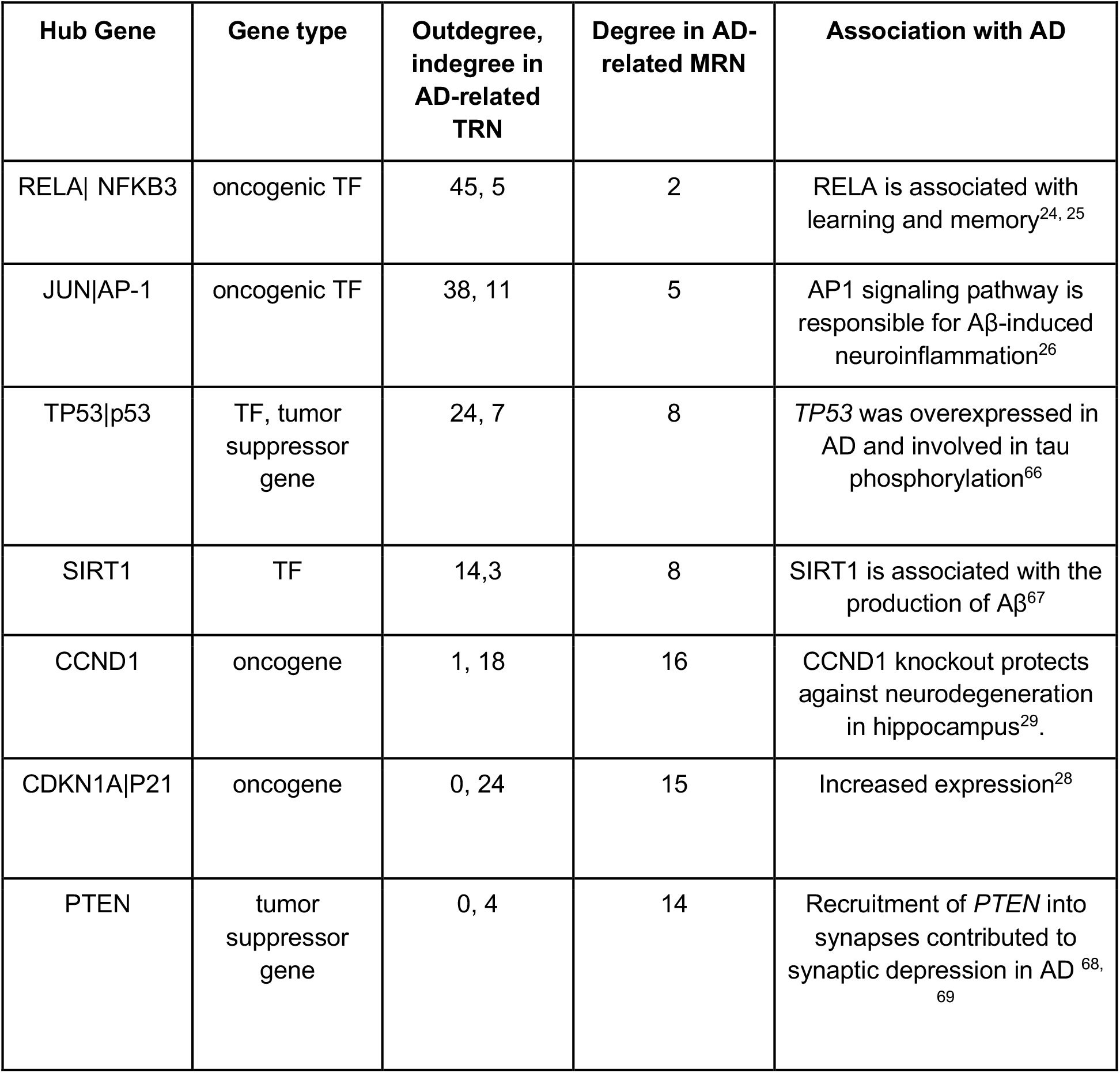
Hub genes (after excluding known AD-associated genes) measured with the outdegree and indegree in AD-related transcriptional regulatory network (TRN) and with the degree in miRNA-based regulatory networks (MRN).

### The top-ranked genes are functionally related to AD based on multiple lines of genomic evidence

#### The top-ranked genes are enriched in AD-associated functions and phenotypes

We hypothesize that genes with higher scores are more likely to be enriched in AD phenotype-related gene sets. To test this hypothesis, we excluded all genes in the training dataset, ranked the remaining ones based on their scores, and tested their enrichment in AD-related gene sets. We collected four gene sets associated with AD pathology. The first gene set was collected from AlzGene, which contained 277 genes. The other three gene sets, namely, the learning or memory pathway (214 genes), the cognition pathway (247 genes), and the amyloid-beta related pathway (51 genes), were collected from the Gene Ontology (GO) database. Using the decile enrichment test (see the *Methods* section), we observed that the top-ranked genes were significantly enriched in the four gene sets: AlzGene (*p-value* = 7.3×10^−13^), learning or memory pathway (*p-value*=6.6×10^−12^), cognition pathway (*p-value* = 1.4×10^−11^), and amyloid-beta pathway (*p-value*=1.1×10^−9^) (Fig. 2B).

We next tested whether the top-ranked genes were functionally similar to AD-associated genes. From the ranked genes, we selected the same number of top-ranked genes as the curated positive genes (n=147). We then performed GO enrichment analysis of both the curated positive genes and the top-ranked genes using PANTHER^19^. The known positive genes and our predicted AD-associated genes were enriched in 771 and 2573 terms, respectively, with 518 of these terms being shared, which was significant compared with the baseline in that no more than 1 pathway was shared (*p-*value<0.01). The 10 most significant shared terms are listed in Supplementary Table 2. We found that many known AD-related functions, including learning or memory, cognition, regulation of endocytosis, regulation of immune system process, regulation of cell death, and regulation of amyloid-beta formation, were shared pathways, implying that our predicted genes might be involved in AD pathology. Specifically, we tested whether the top-scored genes (score > 0.7) were involved in neuron development. Based on GO enrichment analysis, we found that they were enriched in both neuron development (GO:0048666) (FDR = 3.86×10^−76^) and central nervous system neuron development (GO:0021954) (FDR = 5.63×10^−14^).

We further tested whether the top-ranked genes overlap with gene modules that were associated with AD in published studies. A recent study identified gene coexpression modules that were related to AD^5^. Module 109 (m109) containing 390 genes was most strongly associated with cognitive decline. 350 genes overlapped with the brain FGN used in our work and therefore had predicted scores. We found that 101 genes in m109 were among the top-scored genes (score > 0.7), which was significant compared to the random baseline (p < 0.0001). We also obtained two gene sets from another recently published network association study on AD^17^. For protein phosphorylation events in AD, the study derived 28 kinases which were possibly implicated in AD, with 22 kinases having scores >0.7. Among the 14 genes in the amyloid-beta correlated cascade reported by the authors (after removing CLU because it is in the training set), nine had scores > 0.7. These results provide additional evidence that our predicted genes are associated with AD.

#### The top-ranked genes show higher sequence similarity with AD-associated genes

We evaluated whether the sequences of the top-ranked genes were similar to those of AD-associated genes using the sequence similarity method (see the *Methods* section). Let *k*∈[100, 200, 500] denote the number of top-ranked genes for testing. We found that the top-ranked genes had significantly higher sequence similarity with AD-associated genes than randomly selected genes (*p*-value < 0.0001, Supplementary Fig. 6). Taking the top-ranked 200 genes as an example (Fig. 2C), the standardized SEQSIM-score was 6.09, which was significantly higher than that of the randomly selected genes (*SEQSIM*-*score*=-0.0006). The sequence similarity implies the functional similarity between predicted and known AD-associated genes.

#### The top-ranked genes are coexpressed with AD-associated genes

For the top-ranked *k*∈[100, 200, 500] genes, we showed that they were coexpressed with more AD-associated genes than random baseline on the independent Mayo RNA-seq dataset^20^ (*p*-value<0.0001) (Supplementary Fig. 7; see Methods). For example, the number of coexpressed gene pairs between the top-ranked 200 genes and the AD-associated genes was significantly higher than that of randomly selected genes (*p-value* < 0.0001, Fig. 2C), suggesting an association of our top predicted genes with AD.

#### The top-ranked genes interact strongly with AD-associated genes in PPI networks

We hypothesized that the top-ranked *k* genes were more likely to interact with AD-associated genes if the prediction is accurate. We obtained PPI networks from two databases: HuRI and STRING (see Methods). To avoid circularity, we removed those interactions which were used to construct the brain FGN from the two databases. We found that the top-ranked *k*∈[100, 200, 500] genes showed significantly more interactions with AD-associated genes (p-value < 0.0001, Supplementary Fig. 8). Taking the top-ranked 200 genes as an example, the total number of interactions with AD-associated genes was 48 in HuRI, whereas only 11 interactions were found for the randomly selected genes (p-value <0.0001, Fig. 2C).

#### The top-ranked genes are associated with AD based on miRNA-target networks

miRNAs are important post-transcriptional regulators and have been implicated in AD^21^. We investigated whether top-ranked genes were functionally related to AD-associated genes or miRNAs. First, we observed that they shared more miRNAs with AD-associated genes than randomly selected genes (Supplementary Fig. 9; Methods). For instance, the top-ranked 200 genes shared a significant number of miRNAs with AD-associated genes (Fig. 2C, p-value<0.0001). Second, we found that the top-ranked genes interacted with a significant number of AD-associated miRNAs (Fig. 2C; Supplementary Fig. 9). These results imply that top-ranked genes are likely to be involved in post-transcriptional regulatory pathways associated with AD.

### AD-related regulatory networks reveal hub genes and hub miRNAs associated with AD

We constructed two regulatory networks. One is a transcriptional regulatory network (TRN) extracted from the TRRUST database^22^ (version 2.0) that included only known and top-ranked AD-associated genes (Fig. 3A and the *Methods* section). From this network, we identified hub genes based on outdegrees and indegrees. The genes with outdegree and indegree represent transcription factors (TFs) and target genes, respectively. The other regulatory network is a miRNA-target interaction network (Fig. 3B) extracted from mirTarBase^23^ (version 7.0) by considering only AD-associated genes and miRNAs (Methods).

**Fig. 3.**
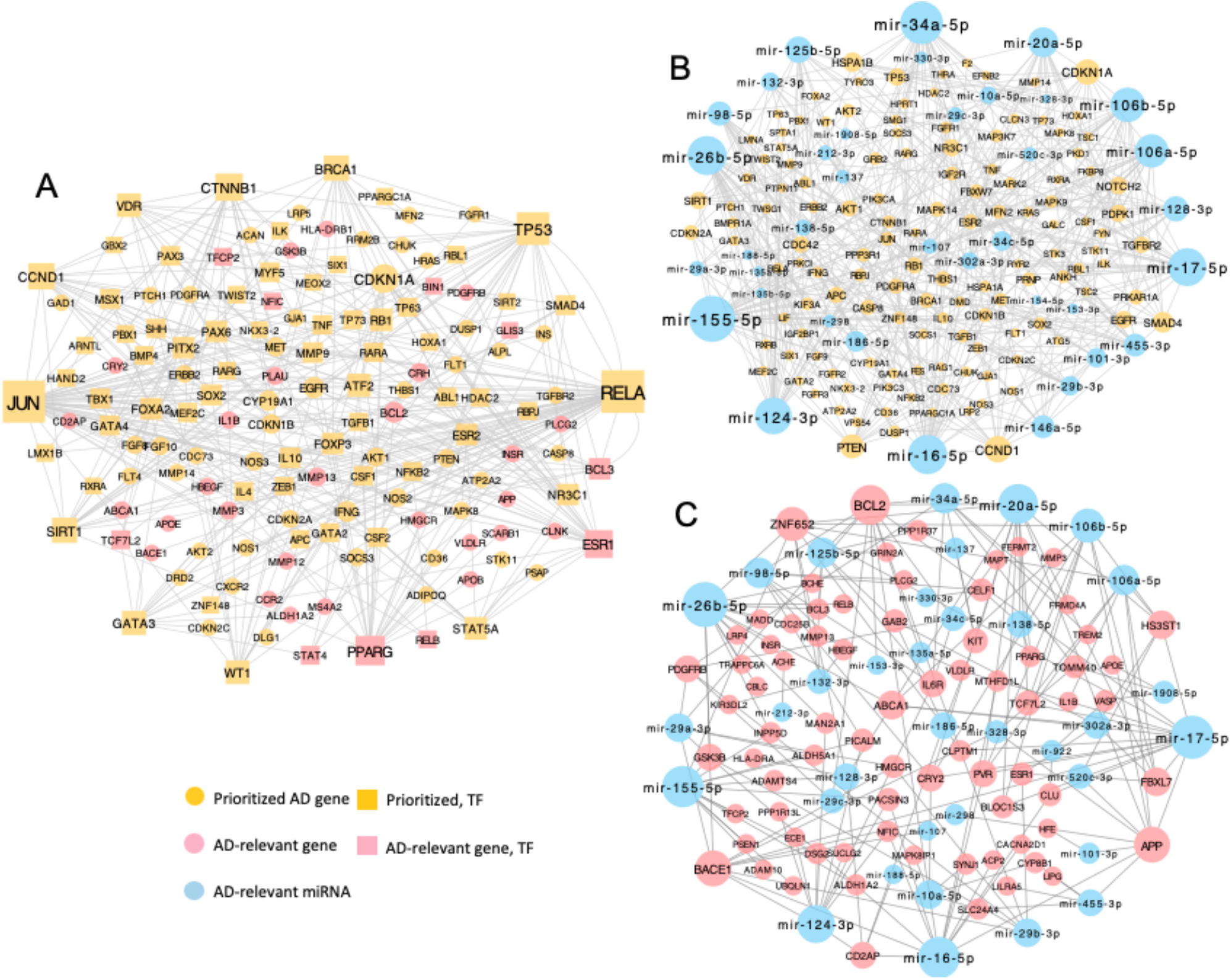
AD-related regulatory networks. **A** Transcriptional regulatory network including our compiled AD-associated genes and the top-ranked genes. **B** The interaction network between predicted genes and AD-relevant miRNAs. **C** The interaction network between the compiled AD-associated genes and AD-relevant miRNAs.

We found that the hub genes in the AD-related TRN were supported by the literature and interaction evidence (Table 1). For example, *RELA* regulates 13 AD-associated genes including *APOE and BACE1*, interacts with 8 AD-associated genes in PPI networks, and is coexpressed with 16 AD-associated genes. Furthermore, *RELA* was shown to be associated with neuroprotection, learning, and memory^24, 25^. Another hub gene is *JUN*. It regulates 11 known AD-associated genes such as *APP, BCL3, RELB*, and *PLAU*, and interacts with the proteins encoded by 10 AD-associated genes such as *MS4A2* and *GSK3B*. Besides, *JUN* is also responsible for Aβ-induced neuroinflammation through a signaling pathway^26^.

We identified genes such as *CCND1* and *CDKN1A* as hubs in the miRNA-based regulatory network (Fig. 3B). Although some studies have reported their associations with AD^27, 28^, the mechanisms underlying these associations are not well understood. These genes might contribute to AD by perturbing the post-transcriptional regulatory network mediated by miRNAs (Table 1 and Fig. 3B). For example, *CCND1* was associated with 16 miRNAs that also bind to known AD-associated genes, including six miRNAs (miR-16-5p, miR-106b-5p, miR-106a-5p, miR-20a-5p, miR-17-5p and miR-101-3p) that bind to *APP* and four miRNAs (miR-29b-3p, miR-186-5p, miR-29c-3p and miR-124-3p) that bind to *BACE1*. In addition, knockout experiments of *CCND1* showed its protective role in neurodegeneration in the hippocampus^29^. Comparing the two networks focusing on only predicted (Fig. 3B) and known (Fig. 3C) AD-associated genes, we observed hub miRNAs such as miR-17b-5p, miR-26b-5p, miR-155-5p, miR-124-3p, and miR-106b-5p that were shared between them, indicating that the shared miRNAs might play roles in the pathology of AD.

### Gene modules in the integrated gene interaction network are associated with AD-related functions, neuropathological and clinical phenotypes in independent data

We constructed an integrated gene interaction network by aggregating multiple lines of genomic evidence and identified four gene modules with a community cluster algorithm (Methods). The modules (denoted by M1, M2, M3, and M4) are shown in Fig. 4 (the genes in each module are provided in Supplementary Table 3). For each module, we performed enrichment analysis using PANTHER^19^ and identified the significantly enriched biological process terms (FDR <0.05). As many of the enriched terms were redundant, we selected representative GO terms with REVIGO^30^. All four modules were enriched in AD-associated biological processes (Fig. 4). For example, M1 was enriched in regulation of cell death and regulation of neurogenesis; M2 was enriched in functions including response to amyloid-beta; M3 was enriched in learning or memory, regulation of synaptic plasticity; M4 was enriched in functions such as regulation of lipid transport and cholesterol efflux. These enrichments imply that the gene modules are not only biologically meaningful but also related to AD.

**Fig. 4.**
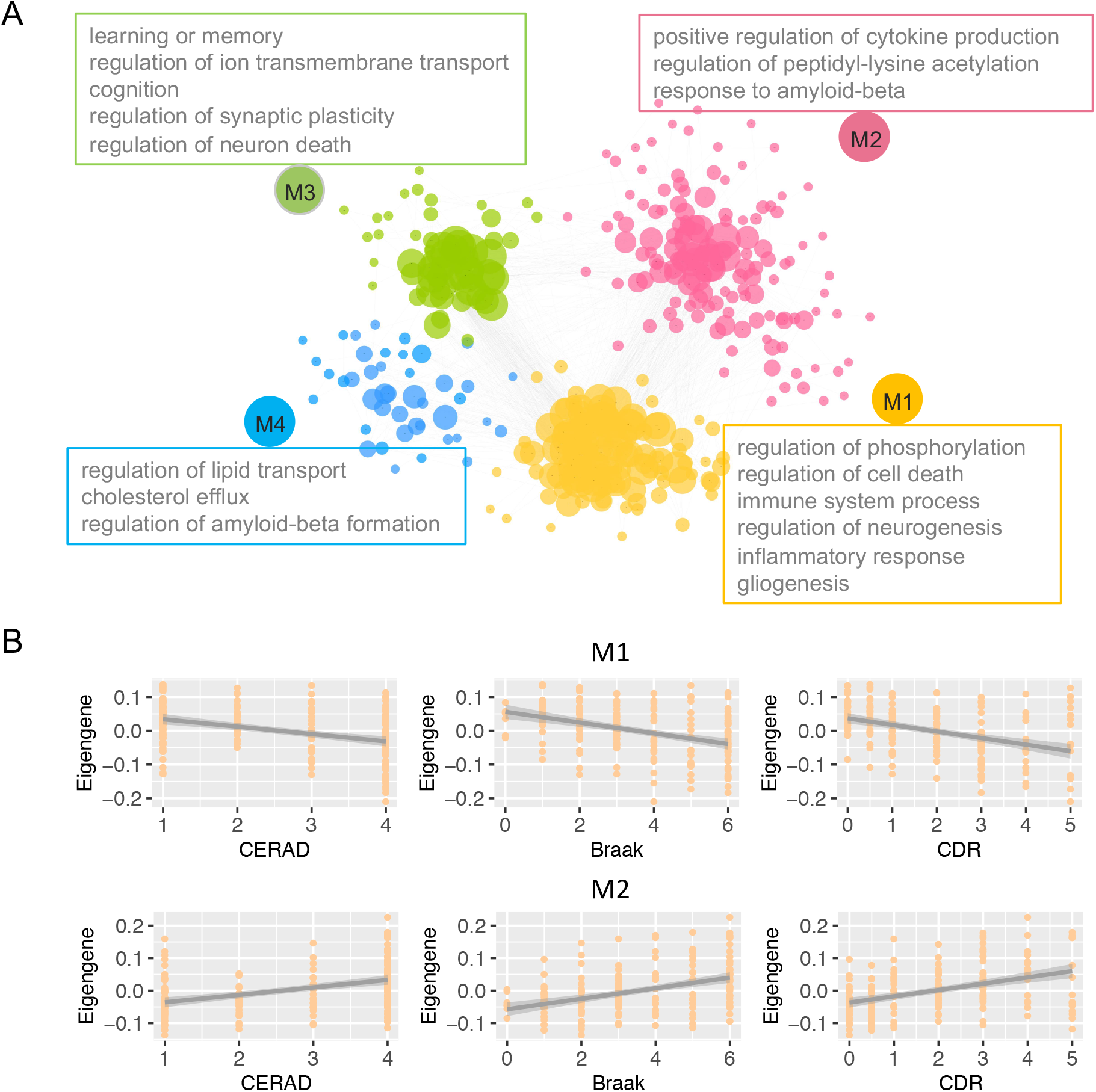
Gene modules and their association with AD traits. The network was built by aggregating the evidence from the protein-protein interaction network, coexpression network, miRNA-gene binding network, transcriptional regulatory network and the brain FGN. This network contains the top-ranked 200 genes and the 147 compiled AD-associated genes. **A** Four gene modules, denoted by M1, M2, M3 and M4, were identified by applying the GLay algorithm to the integrated network in Cytoscape. **B** The association of M1 and M2 with the three AD-related phenotypes (the CERAD, Braak and CDR score) was assessed. The results for all the tests were significant (FDR < 0.05).

Next, we tested whether the modules were correlated with AD-related traits using a well established method^31^. For each module, we extracted the gene expression matrix containing the genes only in that module. We then computed the eigengene (*i*.*e*. the first principal component) of the expression matrix followed by correlating the eigengene with the AD-related traits of interest. We performed this analysis on the independent MSBB RNA-seq dataset with data available for three traits: the CERAD, Braak and CDR score. We conducted a total of twelve correlation tests resulting from all combinations of the four modules and the three traits. We found that the results of all correlation tests were significant (FDR < 0.05), suggesting that our identified modules were associated with AD traits. Taking the eigengene of M1 as an example, it was significantly correlated with the CERAD (r=-0.37, FDR=2.2×10^−7^), Braak (r=-0.41, FDR=1.5×10^−8^), and CDR score (r=-0.42, FDR=6.1×10^−9^) (Figure 4B). Another example was M2, whose eigengene was significantly correlated with the three traits (Figure 4B). The correlation of M3 and M4 with the AD-related traits are provided in Supplementary Fig. 10.

### Individual top-ranked genes are associated with neuropathological and clinical phenotypes on independent datasets

We hypothesized that the top-ranked genes were more likely to be associated with AD-related phenotypes if our prediction was accurate. We tested this hypothesis using the independent MSBB RNA-seq dataset described above. For each gene, we calculated its PCC with the CERAD, Braak and CDR score (see the *Methods* section). To better investigate the trends between our prediction and the gene’s absolute correlation with AD-related phenotypes, we ranked all the predicted genes, divided them into 50 groups, and calculated the mean PCC for each bin. We found that higher ranks (higher predicted scores) were associated with higher mean PCC values for all three phenotypes. The predicted ranks were well correlated with the CERAD (r = 0.68), Braak (r = 0.70) and CDR (r = 0.73) score. The eigengenes for the top-ranked 100, 200 and 500 genes were all significantly correlated with CERAD, Braak and CDR scores (Supplementary Fig. 11).

We then examined the correlations of individual top-ranked genes (those not included in the training set) with AD-related phenotypes^5^. Among the top-ranked 200 genes, we identified 95, 98 and 108 genes that were significantly correlated with CERAD, Braak and CDR scores, respectively (FDR < 0.05). Of them, 84 were correlated with all three phenotypes (Supplementary Table 4). Looking at *FYN*, its correlations with CERAD, Braak and CDR scores were 0.37, 0.35 and 0.37, while *PRKAR1A* had Pearson correlation coefficients of -0.25, -0.31 and -0.29 for the three traits respectively. These results indicate that our top-ranked genes were likely candidate genes for AD.

### Multiple evidence-supported AD-associated genes and their regulatory variants

In the above sections, we have shown that the top-ranked genes are associated with AD based on multiple lines of functional genomic evidence. Here we performed further screening for AD-associated genes by aggregating these evidence, which are divided into two categories: (1) molecular interaction evidence reflecting the interaction of predicted genes with compiled AD-associated genes, and (2) phenotypic correlation evidence supported by correlation of predicted genes with AD traits. The former includes three types of evidence, which are protein interaction, mRNA coexpression, and miRNA sharing with AD-associated genes. The latter includes four types of evidence, which were the correlation with CERAD, Braak and CDR scores based on the MSBB dataset, and differential expression based on the ROSMAP dataset^32^.

To narrow down the predicted candidates, we focused on the top-ranked 200 genes (after excluding the compiled AD-associated genes). The seven types of genomic evidence for these genes are visualized as a circus plot (Figure 5), from which the evidence for each gene can be easily identified. We also obtained their enriched GO biological process terms and showed the functional annotation of these genes (Figure 5). We then applied strict criteria on functional evidence to screen for potentially confident AD-associated genes. That is, only one molecular interaction evidence and one phenotypic correlation evidence is allowed to be missing for each gene. From this, 36 out of the top-ranked 200 genes were retained (Supplementary Table 5), providing a set of multiple evidence-based candidate genes to the community for further functional experiments. As the function of a gene is directly related to the cell type it is expressed in, we further investigated the cell type specificity of their expression. Zhang et al. provides a set of genes that show cell type-specific expression in five major brain cell types including astrocyte, microglia, endothelial, oligodendrocytes and neuron^33^. Using this dataset, we found that 14 of the 36 genes showed specific expression in cell types such as astrocytes and microglia (Supplementary Table 6), while the others are expressed in two or more cell types.

**Fig. 5.**
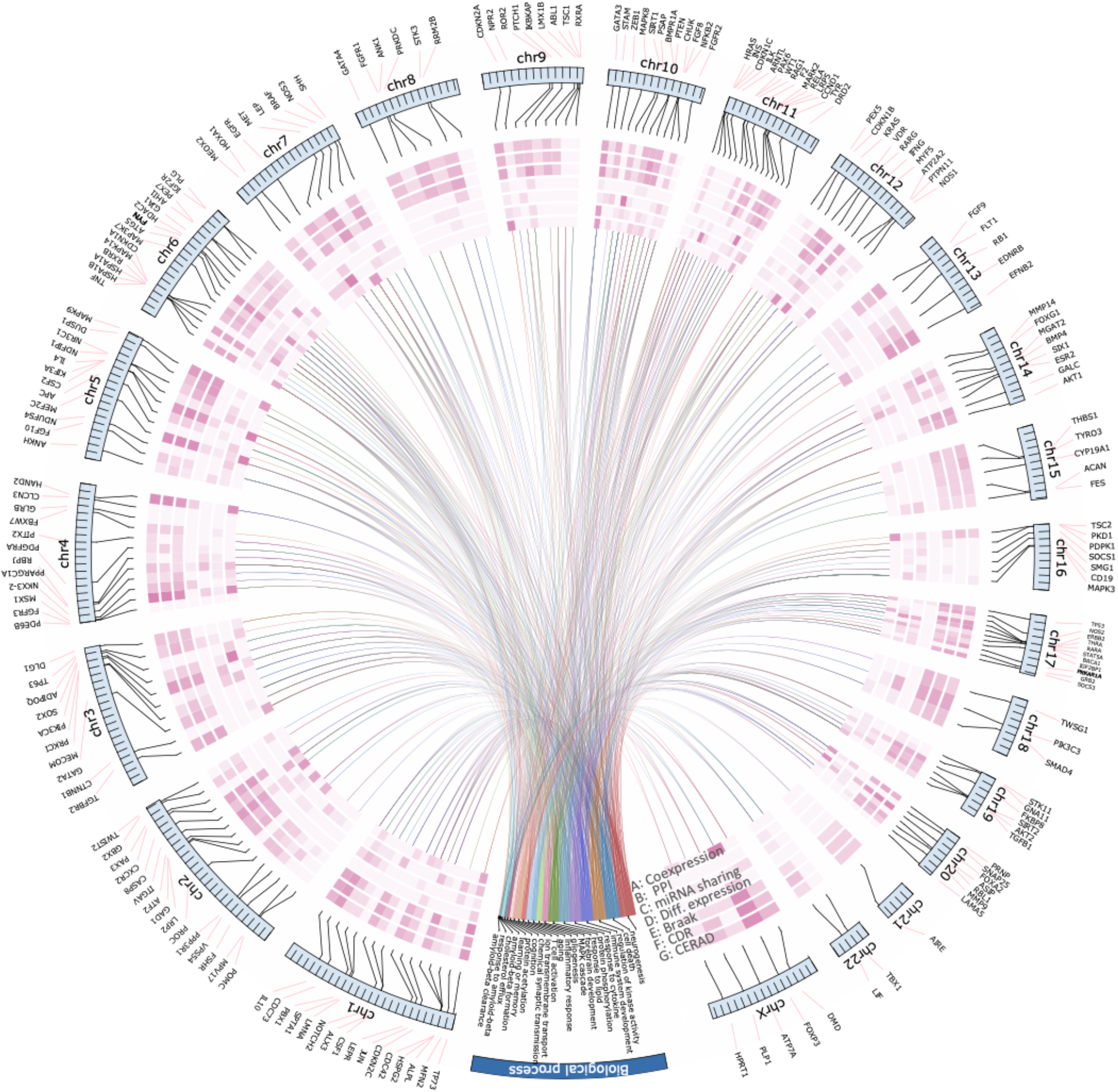
Visualization of functional evidence supporting the association of the top-ranked 200 genes with AD. The seven circles show the strength of the seven types of evidence, including the three molecular interaction evidence (the number of interacting AD-associated genes in PPI, coexpression network and miRNA-target binding network, respectively) and the four phenotypic correlation evidence (the Pearson correlation with CERAD, Braak and CDR on the MSBB dataset, and the log_2_-transformed fold change of expression obtained from the ROSMAP study). The darker the purple color is, the stronger the functional association is. The section corresponding to the blue arc shows the enriched GO biological process terms, where each curve points the gene annotated to the term.

Taking *FYN* as an example, it encodes a membrane-associated tyrosine kinase that is implicated in the control of cell growth and shows specific expression in astrocytes (Supplementary Table 6). It interacts with proteins encoded by 13 AD-associated genes such as *APP* and *MAPT* in PPI, shows significant coexpression with 10 AD-associated genes like *CLU* and interacts with 5 AD-assocaited miRNAs like *hsa-mir-106b*. Its expression was up-regulated based on the ROSMAP dataset (posterior error probability (PEP) =0.04)^32^. Its up-regulation in AD patients was further supported by the positive correlation with CERAD (PCC = 0.37), Braak (PCC = 0.35) and CDR (PCC = 0.37) scores (FDR < 0.001) on the MSBB dataset. The expression of *FYN* for the sample groups partitioned based on CERAD, Braak and CDR scores is shown in Figure 5A. *PRKAR1A* encodes a regulatory subunit of the cAMP-dependent protein kinases involved in the cAMP signaling pathway. It is functionally related with AD-associated genes through PPI, coexpression and miRNA-target network, and its expression is negatively correlated with the above three neuropathological traits (Figure 5A). Altered expression of *PRKAR1A* in AD patients was also identified^34^, providing independent evidence supporting our prediction.

Having shown that the expression level of the above genes was correlated with AD traits, we next exploited which genetic variants (SNP) might causally regulate the expression of these genes by integrating genetic and regulatory data. A SNP is likely causal if it is not only an eQTL but also resides in the transcriptional factor binding site (TFBS) within the promoter of the target gene^34^. By integrating eQTL and ATAC-seq data, we identified seven genes (*FYN, PRKAR1A, PPP3R1, BMPR1A, LMNA, EGFR* and *KRAS*), for which their eQTLs are also located in the TFBS (Supplementary Table 7). For instance, the SNP rs61202914 is an QTL for the expression of a *FYN* isoform. Further, we found that this SNP also resided in the TFBS of multiple transcription factors within the promoter region of *FYN*, thus likely affecting the binding affinity of the transcription factor and therefore expression level. As an illustration, RFX1_HUMAN.H11MO.0.B, which is a motif representing the TFBS of the transcription factor RFX1, harbors the SNP rs61202914 (Figure 6B). This evidence suggests that rs61202914 is likely a variant causally affecting the expression of *FYN*. For *PRKAR1A*, one TFBS in its promoter region harbors its eQTL (rs8080306) (Figure 6B), indicating that rs8080306 is likely a causal variant that regulates the expression of *PRKAR1A*. To summarize, our integrated analysis of eQTL and TFBS in active promoters suggests potential genetic variants that may be associated with AD through regulating the expression of their corresponding target gene. These results may be valuable to prioritize genes for further experimental studies.

**Fig. 6.**
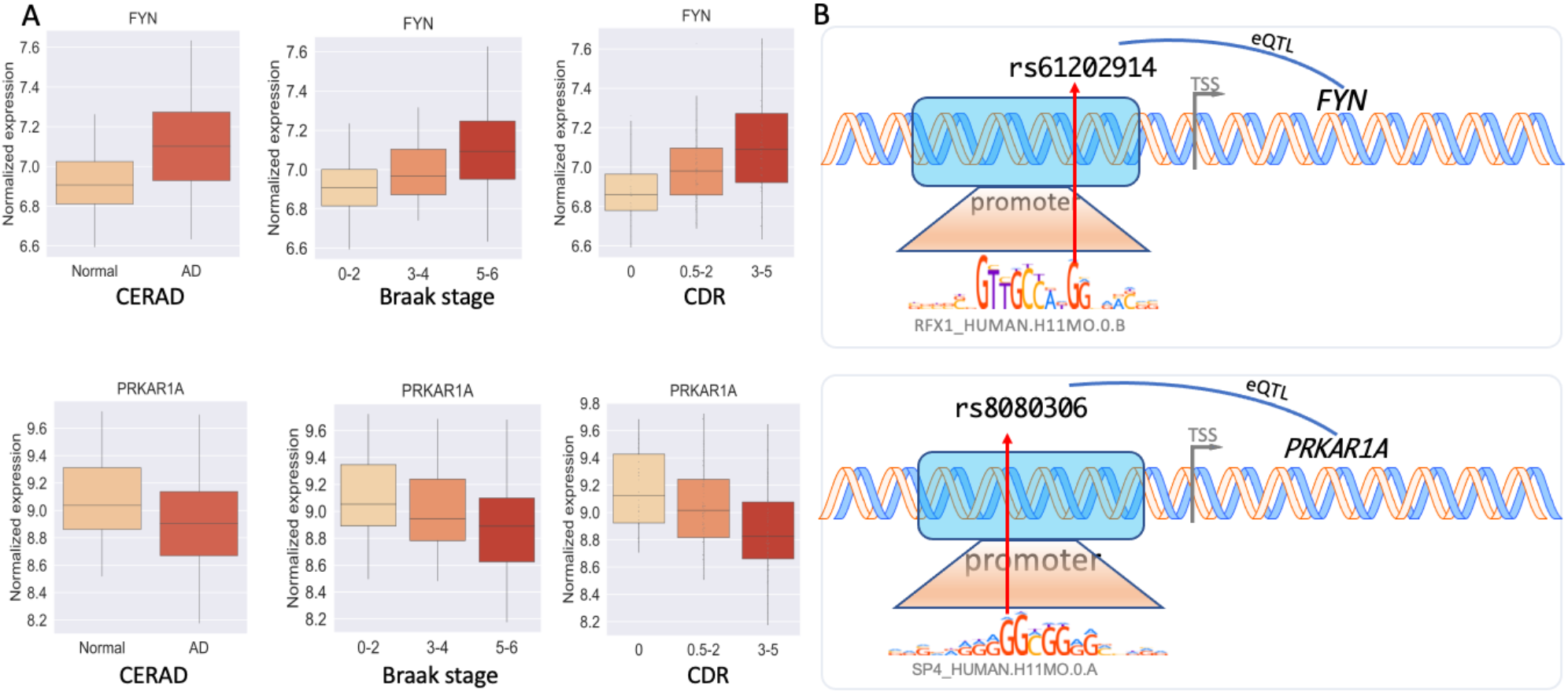
Illustration of the association of the top-ranked individual genes with AD-related phenotypes and the potential regulatory variant of the gene. **A** Comparison of the expression of individual genes in different sample groups. The samples were divided into groups based on the CERAD, Braak or CDR score. The expression for *FYN* and *PRKAR1A* is shown. **B** Potential regulatory SNPs that may regulate the expression. For *FYN*, the SNP rs61202914 not only resides in the TFBS within its promoter region but also is an eQTL (upper); the SNP rs8080306 is located in the TFBS and also an eQTL for *PRKAR1A*.

### ALZLINK: a web resource for interrogating AD-associated genes

To facilitate the interrogation of AD-associated genes and the use of the statistical evaluation pipeline developed in this work, we created the interactive web resource ALZLINK (available at: www.alzlink.com). This site provides the predicted genes along with their predicted scores and functional genomic evidence, facilitating experts in the field of AD to select candidates for further experimental testing. Also, the statistical methods to evaluate the association of an individual gene or a gene set with AD are implemented and available as an online pipeline. For an individual gene, users can query its interactions with known AD-associated genes in heterogeneous interaction networks and its correlation with AD-related traits including CERAD, CDR and Braak scores. For a gene set, users can statistically test its association with AD using the sequence or network-based methods, outputting the distribution of the test metric along with a p-value measuring the significance. For each interaction network such as PPI, the local network involving the queried gene or gene set and the known AD-associated genes is visualized on the web. The data and pipelines on ALZLINK could serve as a valuable resource for experts to prioritize AD-associated genes for further testing.

## Discussion

AD is a neurodegenerative disease with heterogeneous pathologies^8, 35, 36, 37^. However, predicting AD-associated genes is challenging because AD, as a complex disease, is caused mainly by common variants of multiple genes and the disruption of related pathways. FGNs are an important model for characterizing complex functional relationships between genes and have been successfully applied to predict candidate genes for complex diseases, including autism^11^ and Parkinson’s disease^38^. Since AD is caused by gene dysregulation in the brain, we considered brain FGNs as the basis for predicting AD-associated genes. The key idea of our approach was to discover the pattern of AD-associated genes from a brain FGN using machine learning methods. Using our model, we were able to predict novel candidate genes for AD.

We evaluated the association of top-ranked genes with AD by investigating their enrichment in AD-related functions and phenotypes along with examining their association with AD through multiple heterogeneous biological networks. We found that the top-ranked genes were associated with AD. Based on the analyses of the independent MSBB data, we observed that the top-ranked genes were correlated with AD-related neuropathological (CERAD and Braak scores) and clinical (CDR) phenotypes, suggesting that they were likely associated with AD. We also explored gene modules from the AD-related network. We found that these modules were enriched in many AD-related pathways and phenotypes and were also correlated with three AD-related phenotypes, implicating their biological relevance. Combining the genomic data and our predictions, we identified a set of 36 genes whose association with AD was supported by multiple lines of evidence, indicating these genes as potential promising candidates. We further identified potential causal variants for 7 of the 36 genes by integrating brain eQTL and ATAC-seq data.

Our contributions are mainly three-fold. First, we compiled a set of genes that were likely related to AD by performing an intensive, stringent hand curation of multiple resources, providing a potential resource for the community. For negative gene selection, we proposed a pathway-based approach that works by removing any gene that was likely to be associated with AD. Thus, it can be expected that negative genes have been identified. We illustrated that this approach helped improve the accuracies of models in terms of both AUROC and AUPRC. Our model for predicting AD-associated genes depends on the non-AD (negative) genes. Different ways of negative gene selection could lead to bias in the model and thus the prediction. As our method selects negative genes by removing any gene that has a potential association with AD, a possible bias is that the predicted genes are more likely to be functionally related to and share GO terms with the compiled AD-associated genes. Second, we predicted novel candidate genes and showed that the top-ranked genes exhibit significant associations with AD through functional enrichment analysis and the investigation of multiple biological networks. Moreover, the genes were found to be correlated with AD-related phenotypes on independent datasets. Taking advantage of the functional genomic data, we identified a set of 36 AD-associated genes supported by multiple lines of evidence, indicating promising candidates. Third, we developed ALZLINK, a web interface to facilitate the use of data and pipeline developed in this study. It should be pointed out that the pipeline to evaluate the relevance of the predicted genes to AD is generic and can be applied to any other diseases.

Although our predictions are promising, as supported by our systematic analysis, our model for predicting AD-associated genes could be improved in several ways. First, our predictions were made at the gene level without differentiating the splice isoforms generated from the same gene through alternative splicing^39, 40^. This factor is essential because isoforms of the same gene might have different or even opposite functions. Isoforms have been implicated in diseases such as ovarian cancers^41^. The prediction of AD-associated genes at the isoform level could have the potential to promote our understanding of AD. Second, the human brain consists of multiple heterogeneous structures, each of which contains many different cell types. The association of the predicted genes with AD in different cell types remains to be resolved. Integrating single-cell genomic data^42, 43, 44^ with our predicted genes could be helpful for addressing this question. Lastly, our predictions do not implicate causality. The genes predicted using our method are statistically significantly associated with AD.

In summary, we predicted novel AD-associated genes and provided evidence for their association with AD. However, further studies are needed to test the validity of our predictions. This pipeline of prediction and validation is generic and can be readily used for other diseases, such as Parkinson’s disease, cancers and heart diseases. We expect that the predicted genes might become a useful resource for experimental testing by the community and that our proposed pipeline could be used in other diseases.

## Methods

### Compilation of AD-associated and non-AD genes

AD-associated (positives) and non-AD (negatives) genes are needed to build a machine learning model. First, we performed intensive hand-curation to identify confident AD-associated genes from various disease gene resources, including AlzGene^45^, AlzBase^46^, OMIM^47^, DisGenet^48^, DistiLD^49^, and UniProt^50^, Open Targets^51^, GWAS Catalog^52^, differentially expressed genes (DEGs) in ROSMAP^32^ and published literature. The curated genes from each resource as well as the corresponding criteria were provided in Supplementary Note 1. As the AD-associated genes and their reliability vary across these resources, we applied a voting strategy and selected only those that were present in at least two resources to ensure higher reliability (see details in Supplementary Note 1). In this way, we obtained 147 AD-associated genes. Second, we selected a set of non-AD genes, which had no or minimal association with AD. The main idea of our method for non-AD gene selection was to remove any genes that exhibit potential associations with AD. We removed genes that (i) were annotated to the same Gene Ontology (GO) term enriched for the AD-associated genes or (ii) showed any association with AD based on the above-described resources (see details in Supplementary Note 1). In this way, we identified 1651 non-AD genes.

### Model development for predicting AD-associated genes

We first constructed the feature matrix for all human genes based on the brain-specific FGN. This FGN was built by integrating heterogeneous functional genomic data, including gene expression, protein-protein interaction (PPI), protein docking and gene-to-phenotype annotation using the well-established Bayesian framework^16^. The Bayesian network model predicts a co-functional probability (CFP) for every pair of genes by using the following formula:

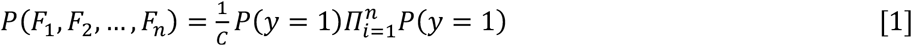

where *P*(*y*=1) is the prior probability for a sample (*i*.*e*. a gene pair in this study) to be positive, *P*(*F*_*i*_|*y* = 1), *i* = 1, 2, …, *n*, is the probability of observing the value of the *i-*th feature under the condition that the gene pair is functionally related, and *C* is a constant normalization factor. In the resulting network, a node is a gene, and an edge represents CFP that two linked genes participate in the same biological process or pathway. For each gene, we extracted its CFP with the compiled AD-associated genes (147 genes) from the network as features based on a previously proposed method^18^. As a result, each gene is characterized by a 147-dimensional vector. The feature data for the training set (147 positives and 1651 negatives, resulting in a total of 1798 genes) are represented by a *1798×147* matrix ***X***. The label (1 for positives and 0 for negatives) of each gene is stored in a vector ***y***. The feature matrix of all other genes not in the training set was extracted.

To develop a model for predicting AD-associated genes, we compared the different combinations of FGNs and machine learning models. To identify optimal FGNs for feature matrix construction, we obtained ten networks for the whole brain or brain-regions, including the brain, forebrain, frontal lobe, temporal lobe, hippocampus, thalamus, amygdala, glia and astrocytes from the GIANT database^15^ and the BaiHui database^16^. We considered these ten regions because they have been implicated in AD^53, 54^. As AD-associated genes are likely to operate in immune cells^55, 56^, we investigated how well immune cells were represented in these networks. As microglia is the dominant immune cell in the brain and cell type-specific genes are indicators of the cell type of interest, we analyzed how microglia-specific genes were represented in these networks. We obtained a set of microglia-specific genes from the work^33^. We found that more than 95% of them existed in each of these networks, suggesting that immune cells are well represented in these networks. For the machine learning models, we considered logistic regression (LR), support vector machine (SVM), random forest (RF) and extremely randomized trees (ExtraTrees) for their promising accuracy shown in our previous work^18^.

### Statistical assessment of the relevance of top-ranked genes to AD

We evaluated the relevance of the top-ranked genes to AD using the following method (the genes in the training set were excluded). These methods are based on the sequence, pathway and various biological networks, as described below.

#### Decile enrichment test for AD pathways and phenotypes

If the prediction is accurate, it is expected that AD-associated genes are more likely to be enriched in the top-ranked genes. Using the decile enrichment test proposed in the previous study^11^, we statistically assessed whether a larger proportion of a given AD-related gene set falls into the first decile of the ranked genes. To do so, we excluded the genes in the training set, ranked the remaining genes, and split genes into 10 evenly binned deciles. Let *P*_*net*_ and *P*_*random*_ denote the proportion of a given gene set that falls into the first decile based on our prediction and random chance, respectively. We tested whether *P*_*net*_ was significantly larger than *P*_*random*_ by using the binomial test (see details in the previous work^11^).

#### Evaluation based on sequence similarity

Genes with similar sequences are likely to carry out similar functions. For a set of *k* predicted genes denoted by *G*_*k*_, we evaluate its functional relationship with AD-associated genes using a sequence similarity-based score (SEQSIM-score), which measures the average similarity between predicted and known AD-associated genes. It is calculated as:

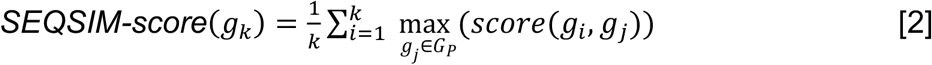

where *G*_*P*_ denotes the set of compiled positive genes, *score(g*_*i*_, *g*_*j*_) is the sequence identity between a predicted gene *g*_*i*_ and the AD-associated gene *g*_*j*_ calculated using BLAST^57^. The higher the *SEQSIM-score* is, the more similar to AD-associated genes the predicted gene is. *SEQSIM-score* was standardized to have zero mean and unit variance using *z*-transform. For the top-ranked *k*∈[100, 200, 500] genes, their scores are denoted by the *SEQSIM-score*_*observed*_. In the same way, we also calculated the *SEQSIM-score*_*random*_ for a set of *k* randomly selected genes. We calculated 10,000 such scores from 10,000 randomly sampled gene sets. Let *N*_*sig*_ denote the number of random scores that are higher than *SEQSIM-score*_*observed*_. We computed the *p*-value as *N*_*sig*_/10000.

#### Evaluation based on coexpression with AD-associated genes

Compared to randomly selected genes, reliably predicted genes are more likely to be coregulated with AD-associated genes. Based on this hypothesis, we calculated the number of coexpressed gene pairs between top-ranked *k* genes and known AD-associated genes using independent gene expression data. That’s to say, in each pair, one is a predicted gene and the other is a known AD-associated gene. The coexpression was measured with Pearson correlation coefficient (PCC). A gene pair was considered to be coexpressed if the PCC ≥ 0.7. To test whether the coexpression is significant, we generated 10,000 gene lists, each containing *k* randomly sampled genes. We calculated the number of coexpressed gene pairs for the top-ranked genes and for the randomly selected genes, denoted by *E*_*observed*_ and *E*_*random*_. We calculated the *p*-value to measure whether *E*_*observed*_ is significantly higher than *E*_*random*_.

We used the Mayo RNA-seq dataset generated from the Accelerating Medicines Partnership-Alzheimer’s Disease (AMP-AD) project (publicly available at https://www.synapse.org/#!Synapse:syn2580853) for coexpression evaluation. Note that this dataset was not used for constructing the brain FGN that was used to build the model for predicting AD-associated genes, so circularity was avoided. This dataset contains gene expression data of the temporal cortex obtained from 82 cases and 80 controls. The log_2_-transformed Fragments Per Kilobase of transcript per Million mapped reads (FPKM) was used for this analysis.

#### Evaluation based on PPI networks

We tested whether the top-ranked *k* genes were more likely to interact with AD-associated genes in PPI networks. We used the PPI data from Human Reference Interactome (HuRI)^58^ and Search Tool for the Retrieval of Interacting Genes/Proteins (STRING)^59^. Because some PPI data were integrated to build the brain FGN, such PPIs have been first removed from the two databases to avoid circularity. The interaction data in HuRI were experimentally identified. In STRING, a score is used to measure the interaction strength between two proteins; a score > 700 indicates an interaction with high confidence. Only the confident interaction was considered. We tested *k* values in [100, 200, 500]. For a given *k* value, we computed the number of genes in the top-ranked *k* genes that interacted with at least one AD-associated gene, denoted by *N*_*observed*_. Similarly, we also calculated *N*_*random*_, which represents the number of genes in *k* randomly sampled genes that interacted with at least one AD-associated gene. With the same method described in the previous section, a *p*-value was calculated to measure the significance.

#### Evaluation based on miRNA-target interaction networks

This analysis was motivated by the assumption that top-ranked genes were more likely related to AD-associated genes or miRNAs based on miRNA-target interaction networks. First, we tested whether top-ranked genes and AD-associated genes share more miRNAs. We downloaded miRNA-target interaction data from miRTarBase^23^, a high-quality database of validated interactions. We computed the number of shared miRNAs of the top-ranked *k*∈[100, 200, 500] genes with AD-associated genes. Based on randomly sampled genes, we calculated a *p*-value to test whether the number of shared miRNAs was significant. Second, we tested top-ranked genes for their binding to AD-associated miRNAs. We retrieved AD-associated miRNAs from the Human microRNA Disease Database (HMDD) (v3.2). Similarly, for the top-ranked *k* genes, we calculated a p-value to measure their significance of binding to AD-associated miRNAs.

### Construction of AD-related regulatory networks

To analyze the regulatory relationship between the predicted candidates and AD-associated genes and obtain hub genes^60, 61^, we constructed two AD-related regulatory networks: one was a transcriptional regulation network, the other was a miRNA-target interaction network.

The human transcriptional regulatory network was downloaded from the Transcriptional Regulatory Relationships Unraveled by Sentence-based Text mining (TRRUST) database^22^. The full network contains 795 transcription factors (TFs) and 2492 target genes. First, we extracted an AD-related transcriptional regulatory network by retaining only the TF-target gene pairs in which one node is known or predicted AD-associated gene (among the top-ranked 200). We identified hub genes according to the outdegree or indegree.

For constructing the AD-related miRNA-target interaction network, we first collected 44 AD-associated miRNAs from an up-to-date review^23^. Then from the above-described miRTarBase^23^ (version 7.0), we extracted two networks. One contains only the interaction between AD-associated miRNAs and AD-associated genes, and the other contains only the interaction between AD-associated miRNAs and predicted AD-associated genes.

### Identification of gene modules in the integrated network

To better understand the functions of the predicted genes, we constructed an integrated network by aggregating evidence from the brain FGN, PPI, coexpression network, miRNA-target network and transcriptional regulatory network. This network included the top-ranked 200 genes and the compiled 147 AD-associated genes. Two genes were connected with an edge if they were direct neighbors in any of the networks above. In detail, all TF-target interactions, which satisfy the above condition, were extracted from the transcriptional regulatory network in the TRRUST database^22^. We also included the genes with a CFP ≥ 0.7, and then expanded the resulting network by including other genes that have a CFP ≥ 0.95 with at least one known AD-associated gene. From the gene coexpression network, we retained only edges with PCCs higher than 0.7. From the PPI network, we included gene pairs whose encoded proteins show interaction in HuRI or STRING. For the miRNA-target interaction data, we computed a network in which the weight of the edge between two genes was calculated as *w*=*N*_*share*_*/N*_*max*_, where *N*_*share*_ represents the number of miRNAs shared by the two genes and *N*_*max*_ =*max*(*N*_*1*_, *N*_*2*_) with *N*_*1*_ and *N*_*2*_ denoting the number of miRNAs binding to the two genes, respectively. The range of *w* is from 0 to 1. The interaction with *w* ≥0.3 was considered. By applying the GLay algorithm implemented in Cytoscape[44] to the integrated network, we identified gene modules within which genes were closely connected.

### The Independent Mountain Sinai Brain Bank (MSBB) dataset with AD-related neuropathological and clinical traits

We obtained an independent dataset with AD-related neuropathological and clinical traits from the MSBB study^62^. We used the data from Brodmann area 36 (parahippocampal gyrus), which is one of the most vulnerable regions to AD^63^. This dataset contains gene expression data from 215 donors for which AD-related phenotypes are also available. These phenotypes include the neuritic plaque density assessed by CERAD score, neurofibrillary tangle severity by Braak score, and severity of dementia by CDR score. The dataset contains 23021 genes measured for the 215 individuals and is available at the AMP-AD portal (https://www.synapse.org/#!Synapse:syn3159438). For each gene, its PCC with the CERAD, Braak and CDR scores was calculated.

Based on the CERAD score, we extracted control and AD samples using the criteria provided on https://www.synapse.org/#!Synapse:syn6101474; based on the Braak score, we followed the practice in ^63^ and divided samples into three groups in the ranges of [0, 2], [3, 4] and [5, 6], representing different levels of tau pathology; Based on CDR, the samples were partitioned into three groups in the range of [0], [0.5, 2] and [3, 5] in the same way as used in ^63^, representing different degrees of severity of clinical dementia.

### Brain eQTL and ATAC-seq data

We identify potentially causal regulatory variants by testing whether eQTL for a target gene also resides in the transcriptional factor binding site (TFBS) in its promoters through the integration of eQTL and ATAC-seq data. Both gene- and isoform-expression eQTLs were considered. We obtained brain gene eQTLs from GTEx (version: v8), PsychEncode (http://resource.psychencode.org/) and the CommonMind Consortium (https://www.synapse.org/#!Synapse:syn4622659). The latter two resources contain isoform eQTLs, which were also used. We used active promoters from the human brain ATAC-seq peak data in the BOCA database^64^. We identified TFBSs in these promoters using the FIMO tool^65^, with the transcription factor binding motif in the HOCOMOCO database (version 11) as reference.

## Data Availability

All accession codes, unique identifiers, or web links for publicly available datasets are described in the paper. All data supporting the findings of the current study are listed in Supplementary Tables 1-7, Supplementary Figures 1-11, and our web interface (www.alzlink.com).

## Code Availability

The codes for model development are publicly available at https://github.com/genemine/alzlink.

## Acknowledgments

This work is supported by the National Key R&D Program of China (No. 2018YFC0910504), the National Natural Science Foundation of China (No. U1909208, 61772552, 61772557), 111 Project (No. B18059), and Hunan Provincial Science and Technology Program (2018WK4001).

The results published here are in part based on data obtained from the AMP-AD Knowledge Portal (https://adknowledgeportal.synapse.org/). The Mayo RNA-seq data were provided by the following sources: The Mayo Clinic Alzheimer’s Disease Genetic Studies, led by Dr. Nilufer Ertekin-Taner and Dr. Steven G. Younkin, Mayo Clinic, Jacksonville, FL using samples from the Mayo Clinic Study of Aging, the Mayo Clinic Alzheimer’s Disease Research Center, and the Mayo Clinic Brain Bank. Data collection was supported through funding by NIA grants P50 AG016574, R01 AG032990, U01 AG046139, R01 AG018023, U01 AG006576, U01 AG006786, R01 AG025711, R01 AG017216, R01 AG003949, NINDS grant R01 NS080820, CurePSP Foundation, and support from Mayo Foundation. Study data includes samples collected through the Sun Health Research Institute Brain and Body Donation Program of Sun City, Arizona. The Brain and Body Donation Program is supported by the National Institute of Neurological Disorders and Stroke (U24 NS072026 National Brain and Tissue Resource for Parkinson’s Disease and Related Disorders), the National Institute on Aging (P30 AG19610 Arizona Alzheimer’s Disease Core Center), the Arizona Department of Health Services (contract 211002, Arizona Alzheimer’s Research Center), the Arizona Biomedical Research Commission (contracts 4001, 0011, 05-901 and 1001 to the Arizona Parkinson’s Disease Consortium) and the Michael J. Fox Foundation for Parkinson’s Research. The MSBB data were generated from postmortem brain tissue collected through the Mount Sinai VA Medical Center Brain Bank and were provided by Dr. Eric Schadt from Mount Sinai School of Medicine.

## Author contributions

C.X.L., H.D.L. and W.S.L. developed the statistical method, performed the analysis, and wrote the manuscript. D.C. and C.X.L developed the web interface. X.M.Z., J.W., F.X.W. and D.W. provided instructions on the analysis. J.X.W. conceived and supervised the research and contributed to the manuscript.

## Additional information

**Supplementary Information** accompanies this paper at http://www.nature.com/ nature communications.

### Competing financial interests

The authors declare no competing financial interests.

## Supplementary information

### Supplementary Notes

Supplementary Note 1. Description for compiling AD-associated genes.

### Supplementary Figures

Supplementary Fig. 1. Comparison in model performance of two methods in negative non-AD gene selection.

Supplementary Fig. 2. Comparison of the negative controls and randomly selected genes based on their association with AD.

Supplementary Fig. 3. Performances of different brain-region networks based on Random Forest (RF).

Supplementary Fig. 4. Performances of different brain-region networks based on support vector machines (SVM).

Supplementary Fig. 5. Performance of different brain-region networks based on logistic regression (LR).

Supplementary Fig. 6. Validation of the top-ranked genes based on sequence similarity with AD-associated genes.

Supplementary Fig. 7. Validation of the top-ranked genes based on their coexpression with known AD-associated genes.

Supplementary Fig. 8. Validation of the top-ranked genes based on protein-protein interaction networks in the STRING and HuRI database.

Supplementary Fig. 9. Validation of the top-ranked genes based on miRNA-target binding networks.

Supplementary Fig. 10. The correlation with three AD traits of the eigengenes of modules 3 and 4.

Supplementary Fig. 11. The correlation with three AD traits of the eigengenes of the top-ranked genes.

### Supplementary Tables

Supplementary Table 1. The top-ranked genes (excluding training set) that are likely associated with AD based on literature.

Supplementary Table 2. The top ten shared GO terms of the 147 AD-associated genes with the top 147 predicted genes.

Supplementary Table 3. Gene modules identified from the integrated gene interaction network.

Supplementary Table 4. The correlation of 84 genes with CERAD, Braak Score and CDR on the MSBB data.

Supplementary Table 5. The seven types of functional evidence for the selected 36 genes.

Supplementary Table 6. The 14 genes with cell type specific expression. Supplementary Table 7. The seven genes with eQTLs located in the transcription factor binding site in the promoter region.

## Notes

### Competing Interest Statement

The authors have declared no competing interest.

