## Supplementary Tables for "Genome-wide prediction and integrative functional characterization of Alzheimer’s disease-associated genes"

### List of Supplementary Tables

| Tables | Description |
|---|---|
| Supplementary Table 1 | The top-ranked genes (excluding training set) that are likely associated with AD based on literature. |
| Supplementary Table 2 | The top ten shared GO terms of the 147 AD-associated genes with the top 147 predicted genes. |
| Supplementary Table 3 | Gene modules identified from the integrated gene interaction network. |
| Supplementary Table 4 | The correlation of 84 genes with CERAD, Braak Score and CDR on the MSBB data. |
| Supplementary Table 5 | The seven types of functional evidence for the selected 36 genes. |
| Supplementary Table 6 | The 14 genes with cell type specific expression. |
| Supplementary Table 7 | The seven genes with eQTLs located in the transcription factor binding site in the promoter region. |

Supplementary Table 1. The top-ranked genes (excluding training set) that are likely associated with AD based on literature. (Literature survey was conducted for the top-ranked 20 genes; those with literature evidence are listed below)

| **Gene** | **Rank** | **Description** | **References** |
|---|---|---|---|
| PRKAR1A | 1 | It is a dysregulated gene in AD and a top highly connected gene in immune response. | Canchi S, et al. Integrating Gene and Protein Expression Reveals Perturbed Functional Networks in Alzheimer's Disease. Cell Reports 28, 1103-1116.e1104 (2019). |
| NOS3 | 2 | The nitric oxide synthase 3 G894T polymorphism is associated with Alzheimer's disease risk: a meta-analysis | Liu S, Zeng F, Wang C, Chen Z, Zhao B, Li K. The nitric oxide synthase 3 G894T polymorphism associated with Alzheimer's disease risk: a meta-analysis. *Scientific reports* 5, 13598-13598 (2015). |
| MAPK8 | 3 | JNK1 deficiency resulted in reduced BACE1 expression, suggesting alterations in amyloidogenic pathway. | Petrov D*, et al.* Evaluation of the Role of JNK1 in the Hippocampus in an Experimental Model of Familial Alzheimer's Disease. *Molecular Neurobiology* 53, 6183-6193 (2016). |
| ABL1 | 4 | It is activated in human Alzheimer's. | Schlatterer SD, Acker CM, Davies P. c-Abl in neurodegenerative disease. *J Mol Neurosci* 45, 445-452 (2011). |
| PRKCI | 5 | It activates β-secretase and increases Aβ 1-40/42 and phospho-tau in mouse brain and isolated neuronal cells. | Sajan MP*, et al.* Atypical PKC, PKCλ/ι, activates β-secretase and increases Aβ(1-40/42) and phospho-tau in mouse brain and isolated neuronal cells, and may link hyperinsulinemia and other aPKC activators to development of pathological and memory abnormalities in Alzheimer's disease. *Neurobiol Aging* 61, 225-237 (2018). |
| MECOM | 9 | A GWAS found that this gene is associated with AD with suggestive evidence (pvalue: 8E-7). | Mez J*, et al.* Two novel loci, COBL and SLC10A2, for Alzheimer's disease in African Americans. *Alzheimers Dement* 13, 119-129 (2017). |

| | | | |
|---|---|---|---|
| ATG5 | 10 | Plasma ATG5 is increased in Alzheimer’s disease. | Cho S-J, Lim HJ, Jo C, Park MH, Han C, Koh YH. Plasma ATG5 is increased in Alzheimer’s disease. *Scientific Reports* 9, 4741 (2019). |
| PDPK1 | 12 | It decreases TACE-mediated α-secretase activity and promotes disease progression in prion and Alzheimer's diseases. | Pietri M, et al. PDK1 decreases TACE-mediated α-secretase activity and promotes disease progression in prion and Alzheimer's diseases. Nature Medicine 19, 1124-1131 (2013). |
| LEP | 14 | Leptin dysfunction is associated with Alzheimer's Disease. | McGuire MJ, Ishii M. Leptin Dysfunction and Alzheimer's Disease: Evidence from Cellular, Animal, and Human Studies. *Cell Mol Neurobiol* 36, 203-217 (2016). |
| EGFR | 15 | EGFR is a preferred target for treating Amyloid-β–induced memory loss. | Wang L*, et al.* Epidermal growth factor receptor is a preferred target for treating Amyloid-β–induced memory loss. *Proceedings of the National Academy of Sciences* 109, 16743 (2012). |
| PRNP | 19 | A nonsense mutation of this gene is associated with AD. | Rita Guerreiro et al., A nonsense mutation in PRNP associated with clinical Alzheimer's disease. Neurobiol Aging. 2014, 35(11): 2656.e13–2656.e16 |
| BMPR1A | 20 | Three SNPs of BMPR1A are associated with LOAD in a Chr10-specific association study. | Grupe A*, et al.* A Scan of Chromosome 10 Identifies a Novel Locus Showing Strong Association with Late-Onset Alzheimer Disease. *The American Journal of Human Genetics* 78, 78-88 (2006). |

Supplementary Table 2. The top ten shared GO terms of the 147 AD-associated genes with top 147 predicted genes.

| **GO ID** | **GO terms** | **FDR for known genes** | **FDR for predicted genes** |
|---|---|---|---|
| GO:0043269 | regulation of ion transport | 1.56E-10 | 1.82E-06 |
| GO:0030100 | regulation of endocytosis | 7.03E-09 | 8.96E-03 |
| GO:1902003 | regulation of amyloid-beta formation | 1.03E-08 | 1.60E-02 |
| GO:0007611 | learning or memory | 1.72E-08 | 4.77E-04 |
| GO:0050890 | cognition | 1.76E-08 | 7.02E-05 |
| GO:1905952 | regulation of lipid localization | 8.95E-08 | 7.15E-07 |
| GO:0042327 | positive regulation of phosphorylation | 1.43E-07 | 3.39E-32 |
| GO:0002682 | regulation of immune system process | 2.19E-07 | 1.56E-25 |
| GO:0010941 | regulation of cell death | 4.53E-07 | 1.14E-43 |
| GO:0050776 | regulation of immune response | 1.18E-06 | 2.03E-10 |

Supplementary Table 3. Gene modules identified from the integrated gene interaction network.

| **M1** | **M2** | **M3** | **M4** |
|---|---|---|---|
| CDKN1A | SIX1 | PBX1 | TBX1 |
| TNF | MYF5 | MEF2C | FGF10 |
| EGFR | WT1 | ATF2 | ARNTL |
| CSF1 | HDAC2 | STAT4 | BIN1 |
| PDGFRB | NOS2 | DUSP1 | MS4A2 |
| TGFB1 | SIRT1 | PTCH1 | FGF8 |
| MET | MMP13 | GAD1 | CRY2 |
| ABL1 | CTNNB1 | HMGCR | LRP5 |
| SOCS3 | RB1 | ATP2A2 | GAB2 |
| ILK | BCL3 | GSK3B | TSC2 |
| ERBB2 | RELA | APP | TSC1 |
| TGFBR2 | IFNG | MAPK9 | RIN3 |
| FLT4 | GATA2 | PRNP | ABI3 |
| PTEN | PPARG | NDFIP1 | VASP |
| CDKN2A | CYP19A1 | MAPK8IP1 | BCAM |
| PLAU | TP53 | PPP1R13L | LAMA5 |
| AKT2 | ESR1 | PRKCI | APOC1 |
| INSR | BRCA1 | MAPT | SYK |
| NFKB2 | ZEB1 | MARK4 | POMC |
| MAPK8 | CCND1 | GRIN2A | BCHE |
| PDGFRA | PAX6 | PTK2B | ABCA7 |
| RBPJ | NOS1 | FBXW7 | CD33 |
| AKT1 | NR3C1 | UBQLN1 | CHRNA7 |
| STK11 | JUN | PSEN2 | ACE |
| FLT1 | CSF2 | ATG5 | APH1B |
| PLCG2 | TCF7L2 | CNTNAP2 | NME8 |
| MMP14 | GATA3 | PSMA1 | PVRL2 |
| FGFR1 | PAX3 | APOC4 | APOC2 |
| BMP4 | FOXA2 | ZCWPW1 | |
| CD2AP | CDC73 | SYNJ1 | |
| CHUK | TP73 | CACNA2D1 | |
| CDKN2C | ESR2 | KRAS | |
| THBS1 | MMP9 | AHI1 | |
| ACAN | FOXP3 | ANK1 | |
| GRB2 | HBEGF | HR | |
| BMPR1A | VDR | MADD | |
| PTPN11 | MMP3 | HPRT1 | |
| LEPR | IL4 | PACSIN3 | |

| | | |
|---|---|---|
| FYN | IL1B | ACP2 |
| KIT | RARG | VPS54 |
| PDPK1 | PITX2 | NDUFS4 |
| FGFR2 | GATA4 | PFDN1 |
| NOTCH2 | RXRA | ADAM10 |
| MAPK14 | ABCA1 | ANKH |
| MARK2 | CRH | CACNA1G |
| CDC42 | ZNF148 | RBFOX1 |
| PLG | ADIPOQ | CHRNB2 |
| HSPA1A | TFCP2 | VSNL1 |
| HSPA1B | CDKN1B | SCN8A |
| SOCS1 | MSX1 | FGF9 |
| BRAF | SOX2 | SORCS1 |
| MAP3K7 | NKX3-2 | GLRB |
| STK3 | SMAD4 | KIF3A |
| TYRO3 | BCL2 | SNAP25 |
| FSHR | CCR2 | BZW2 |
| CDKN1C | TP63 | SCN1A |
| LIPG | RELB | DPP10 |
| DMD | STAT5A | PPP3R1 |
| FGFR3 | IL10 | CLASRP |
| ROR2 | RBL1 | PRKAR1A |
| DDR2 | RARA | STAM |
| AMHR2 | CLNK | |
| RET | NOS3 | |
| IGF1R | GLIS3 | |
| IKBKB | INS | |
| ACVR1 | CXCR2 | |
| PTPN6 | APC | |
| FCGR2B | SHH | |
| MYLK | PPARGC1A | |
| INPP5D | TWIST2 | |
| MAPK3 | LMX1B | |
| ADRBK1 | HAND2 | |
| CSF1R | HOXA1 | |
| PTK2 | CD36 | |
| RAF1 | RRM2B | |
| GNA11 | NFIC | |
| NTRK2 | MMP12 | |
| RAC1 | BACE1 | |
| GNAQ | GJA1 | |
| SIK3 | GBX2 | |

| | |
|---|---|
| PKD1 | APOB |
| ACVR2B | DRD2 |
| CDK5 | CASP8 |
| DDR1 | VLDLR |
| GNAO1 | SIRT2 |
| CDK4 | APOE |
| NPR2 | MEOX2 |
| ERN1 | MFN2 |
| KDR | DLG1 |
| AXL | ALPL |
| STAT3 | HRAS |
| FES | ALDH1A2 |
| CDK2 | RXRB |
| ACVRL1 | UBE2L3 |
| IL6ST | PSEN1 |
| HCK | CLU |
| CAMK4 | CDC25B |
| PTPRS | LRP2 |
| EPHB6 | FKBP8 |
| MERTK | CBLC |
| L1CAM | PIK3CA |
| ERBB3 | THRA |
| IL1R1 | TRAPPC6A |
| MUSK | PVR |
| NOTCH3 | SCIMP |
| TNFRSF1A | SPON1 |
| IKBKAP | MS4A4A |
| CD19 | PLP1 |
| HSPG2 | TWSG1 |
| CD44 | FOXH1 |
| NOTCH1 | PRLR |
| RAB27A | LEP |
| SELP | NFATC4 |
| ITGB3 | LIF |
| TLR4 | CREBBP |
| ST14 | IL4R |
| F2 | AR |
| ACVR2A | PITX3 |
| TNFRSF1B | PAX1 |
| MYD88 | TYR |
| HFE | RUNX1 |
| FAS | LRP4 |

| | |
|---|---|
| ICAM1 | SUCLG2 |
| IL6 | CCRL2 |
| TLR2 | FBXL7 |
| CCL13 | HS3ST1 |
| HLA-DQB1 | SLC24A4 |
| LRRK2 | CHRNA2 |
| TIE1 | BLOC1S3 |
| NCF1 | CLCN3 |
| LTA | FERMT2 |
| NRP2 | ITGAV |
| COL1A1 | PICALM |
| LMNA | MAN2A1 |
| GUCY2C | ECE1 |
| EFNB2 | CLMN |
| IL6R | HARBI1 |
| EDNRB | ZNF652 |
| | PPP1R37 |
| | ADAMTS4 |

Supplementary Table 4. The correlation of 84 genes with CERAD, Braak Score and CDR on the MSBB data.

| Gene | CERAD | Braak Score | CDR |
|---|---|---|---|
| PRKAR1A | -0.2494 | -0.3104 | -0.2936 |
| NOS3 | 0.1786 | 0.224 | 0.2054 |
| MAPK8 | -0.3196 | -0.3768 | -0.3583 |
| ABL1 | 0.3289 | 0.331 | 0.2981 |
| PRKCI | -0.282 | -0.3211 | -0.3025 |
| MECOM | 0.2073 | 0.1908 | 0.298 |
| ATG5 | -0.2869 | -0.3134 | -0.332 |
| CSF1 | 0.3763 | 0.4055 | 0.3244 |
| PDPK1 | -0.1995 | -0.1682 | -0.2289 |
| EGFR | 0.2545 | 0.2687 | 0.238 |
| PRNP | -0.2775 | -0.2522 | -0.3323 |
| BMPR1A | 0.2749 | 0.238 | 0.22 |
| CDKN2A | 0.2111 | 0.192 | 0.213 |
| EFNB2 | -0.307 | -0.2404 | -0.2237 |
| DMD | -0.2674 | -0.178 | -0.219 |
| KIF3A | -0.4138 | -0.376 | -0.4195 |
| PIK3C3 | -0.1753 | -0.25 | -0.2247 |
| PTEN | -0.1685 | -0.2189 | -0.2302 |
| RXRA | 0.2453 | 0.348 | 0.2502 |
| NOS2 | -0.2788 | -0.2525 | -0.3434 |
| SNAP25 | -0.4178 | -0.3794 | -0.4321 |
| RARG | 0.3109 | 0.3332 | 0.2517 |
| MAPK9 | -0.4069 | -0.3992 | -0.4467 |
| SIX1 | 0.218 | 0.2187 | 0.3446 |
| FBXW7 | -0.4564 | -0.4054 | -0.4809 |
| MSX1 | 0.2322 | 0.2573 | 0.2564 |
| GJA1 | 0.3089 | 0.3321 | 0.3049 |
| ATP2A2 | -0.3051 | -0.2898 | -0.3616 |
| TGFBR2 | 0.356 | 0.3352 | 0.3404 |
| STK3 | 0.2373 | 0.1839 | 0.2422 |
| HPRT1 | -0.4036 | -0.4089 | -0.4325 |
| STAT5A | 0.3889 | 0.3888 | 0.311 |
| TP73 | 0.2965 | 0.3542 | 0.2747 |
| MAPK14 | -0.2021 | -0.1813 | -0.2117 |

| | | | |
|---|---|---|---|
| RBL1 | 0.2646 | 0.2879 | 0.2893 |
| NDFIP1 | -0.3704 | -0.3621 | -0.4168 |
| FGFR1 | 0.2705 | 0.2751 | 0.2844 |
| TP53 | 0.3548 | 0.2826 | 0.2981 |
| ACAN | 0.2169 | 0.2158 | 0.1886 |
| CYP19A1 | 0.2574 | 0.1757 | 0.1631 |
| IGF2R | 0.2775 | 0.2105 | 0.2818 |
| CDKN2C | 0.2027 | 0.1933 | 0.1749 |
| BRAF | -0.3444 | -0.3838 | -0.3459 |
| GRB2 | -0.2274 | -0.2166 | -0.1755 |
| LMNA | 0.2628 | 0.2533 | 0.2843 |
| KRAS | -0.3282 | -0.353 | -0.3265 |
| FGFR2 | 0.263 | 0.2393 | 0.2244 |
| PTCH1 | 0.2394 | 0.2865 | 0.2373 |
| BRCA1 | 0.3312 | 0.2379 | 0.2638 |
| STAM | -0.2657 | -0.3296 | -0.3566 |
| FES | 0.1728 | 0.1716 | 0.1913 |
| NFKB2 | 0.2881 | 0.3403 | 0.277 |
| SMAD4 | 0.3475 | 0.3314 | 0.3444 |
| MET | -0.2153 | -0.2578 | -0.3393 |
| LRP5 | 0.2857 | 0.3207 | 0.2711 |
| HSPG2 | 0.2729 | 0.2621 | 0.309 |
| DUSP1 | 0.1603 | 0.2714 | 0.284 |
| NDUFS4 | -0.2483 | -0.3519 | -0.3104 |
| ATF2 | -0.2738 | -0.2794 | -0.2807 |
| ILK | 0.1835 | 0.1811 | 0.2671 |
| CCND1 | 0.2306 | 0.1827 | 0.2701 |
| CASP8 | 0.2705 | 0.284 | 0.2084 |
| PRKDC | -0.2018 | -0.3113 | -0.2896 |
| TWSG1 | 0.227 | 0.17 | 0.211 |
| TGFB1 | 0.3167 | 0.318 | 0.3003 |
| FYN | 0.3714 | 0.3518 | 0.3654 |
| MMP14 | 0.3526 | 0.283 | 0.3119 |
| NOTCH2 | 0.4017 | 0.3863 | 0.3891 |
| PAX6 | 0.3007 | 0.2709 | 0.3066 |
| ERBB2 | 0.3394 | 0.2842 | 0.3422 |
| FKBP8 | -0.2344 | -0.2556 | -0.2492 |
| RELA | 0.2718 | 0.3455 | 0.2754 |
| LAMA5 | 0.1925 | 0.2927 | 0.2367 |

| | | | |
|---|---|---|---|
| PPP3R1 | -0.3697 | -0.3286 | -0.3947 |
| LRP2 | 0.2135 | 0.1886 | 0.206 |
| SOX2 | 0.2016 | 0.2102 | 0.2656 |
| MFN2 | -0.2857 | -0.3028 | -0.3023 |
| GAD1 | -0.3238 | -0.3841 | -0.3292 |
| GATA2 | 0.2482 | 0.2453 | 0.2284 |
| FGF9 | -0.3549 | -0.369 | -0.3517 |
| AKT2 | 0.2348 | 0.2982 | 0.2613 |
| MEF2C | -0.3742 | -0.3875 | -0.4007 |
| GLRB | -0.4088 | -0.3866 | -0.4362 |
| FOXG1 | -0.2719 | -0.1794 | -0.2977 |

Supplementary Table 5. The seven types of functional evidences for the selected 36 genes. PEP: posterior error probability; FDR: false discovery rate. (*Note: Meaning of NA: for the 'log2.AD.NDC' column, the data are available for only those significantly differentially expressed genes from the ROSMAP paper (Canchi et al.,Cell Rep, 2019(28)1103-1116); the data for other genes are not available and are thus denoted by NA*)

| Genes | Predicted_scores | miRNA | PPI | coexp | CERAD | FDR (CERAD) | CDR | FDR (CDR) | Braak | FDR (Braak) | log2.AD.NDC | PEP (log2.AD.NDC) |
|---|---|---|---|---|---|---|---|---|---|---|---|---|
| PRKAR1A | 0.97172 | 7 | 1 | 15 | -0.2494 | 0.0016 | -0.2936 | 0.0002 | -0.3104 | 0.0001 | -0.2093 | 0.0956 |
| MAPK8 | 0.97134 | 4 | 12 | 24 | -0.3196 | 0 | -0.3583 | 0 | -0.3768 | 0 | -0.2221 | 0.0004 |
| ABL1 | 0.96910 | 7 | 6 | 26 | 0.3289 | 0 | 0.2981 | 0.0001 | 0.331 | 0.0001 | 0.2123 | 0.0008 |
| PRKCI | 0.96883 | 3 | 1 | 19 | -0.282 | 0.0003 | -0.3025 | 0.0001 | -0.3211 | 0.0001 | -0.1345 | 0.1319 |
| CSF1 | 0.96622 | 4 | 7 | 19 | 0.3763 | 0 | 0.3244 | 0 | 0.4055 | 0 | 0.3527 | 0.0333 |
| EGFR | 0.96476 | 14 | 17 | 7 | 0.2545 | 0.0013 | 0.238 | 0.0026 | 0.2687 | 0.0011 | NA | NA |
| PRNP | 0.96464 | 13 | 6 | 23 | -0.2775 | 0.0004 | -0.3323 | 0 | -0.2522 | 0.0023 | NA | NA |
| BMPR1A | 0.96459 | 12 | 1 | 7 | 0.2749 | 0.0005 | 0.22 | 0.0055 | 0.238 | 0.0041 | NA | NA |
| KIF3A | 0.96282 | 9 | 3 | 25 | -0.4138 | 0 | -0.4195 | 0 | -0.376 | 0 | NA | NA |
| RXRA | 0.96164 | 2 | 4 | 9 | 0.2453 | 0.002 | 0.2502 | 0.0015 | 0.348 | 0 | 0.2576 | 0.0039 |
| SNAP25 | 0.96144 | 5 | 3 | 27 | -0.4178 | 0 | -0.4321 | 0 | -0.3794 | 0 | -0.4932 | 0.0204 |
| MAPK9 | 0.96045 | 7 | 5 | 29 | -0.4069 | 0 | -0.4467 | 0 | -0.3992 | 0 | -0.2449 | 0.0935 |
| FBXW7 | 0.95917 | 12 | 3 | 25 | -0.4564 | 0 | -0.4809 | 0 | -0.4054 | 0 | NA | NA |
| GJA1 | 0.95876 | 6 | 0 | 8 | 0.3089 | 0.0001 | 0.3049 | 0.0001 | 0.3321 | 0 | 0.4554 | 0.0339 |
| ATP2A2 | 0.95852 | 11 | 1 | 26 | -0.3051 | 0.0001 | -0.3616 | 0 | -0.2898 | 0.0004 | -0.2460 | 0.0067 |
| TGFBR2 | 0.95816 | 17 | 1 | 7 | 0.356 | 0 | 0.3404 | 0 | 0.3352 | 0 | NA | NA |
| HPRT1 | 0.95684 | 5 | 0 | 28 | -0.4036 | 0 | -0.4325 | 0 | -0.4089 | 0 | -0.4105 | 0.0019 |
| STAT5A | 0.95673 | 3 | 6 | 16 | 0.3889 | 0 | 0.311 | 0.0001 | 0.3888 | 0 | NA | NA |
| RBL1 | 0.95612 | 7 | 0 | 4 | 0.2646 | 0.0008 | 0.2893 | 0.0002 | 0.2879 | 0.0004 | NA | NA |
| TP53 | 0.95536 | 21 | 12 | 0 | 0.3548 | 0 | 0.2981 | 0.0001 | 0.2826 | 0.0006 | NA | NA |
| LMNA | 0.95343 | 4 | 0 | 10 | 0.2628 | 0.0009 | 0.2843 | 0.0003 | 0.2533 | 0.0022 | NA | NA |
| KRAS | 0.95251 | 15 | 5 | 27 | -0.3282 | 0 | -0.3265 | 0 | -0.353 | 0 | -0.1735 | 0.0166 |
| BRCA1 | 0.95157 | 11 | 4 | 0 | 0.3312 | 0 | 0.2638 | 0.0008 | 0.2379 | 0.0041 | NA | NA |
| STAM | 0.95151 | 1 | 5 | 7 | -0.2657 | 0.0008 | -0.3566 | 0 | -0.3296 | 0.0001 | -0.1782 | 0.2301 |
| NFKB2 | 0.95118 | 3 | 5 | 12 | 0.2881 | 0.0002 | 0.277 | 0.0004 | 0.3403 | 0 | NA | NA |
| MET | 0.95091 | 15 | 3 | 2 | -0.2153 | 0.0071 | -0.3393 | 0 | -0.2578 | 0.0018 | NA | NA |
| CASP8 | 0.94906 | 9 | 4 | 0 | 0.2705 | 0.0006 | 0.2084 | 0.0088 | 0.284 | 0.0005 | NA | NA |
| TGFB1 | 0.94862 | 10 | 4 | 9 | 0.3167 | 0.0001 | 0.3003 | 0.0001 | 0.318 | 0.0001 | 0.3375 | 0.0366 |
| FYN | 0.94858 | 5 | 19 | 10 | 0.3714 | 0 | 0.3654 | 0 | 0.3518 | 0 | 0.1753 | 0.0416 |
| NOTCH2 | 0.94848 | 16 | 5 | 9 | 0.4017 | 0 | 0.3891 | 0 | 0.3863 | 0 | NA | NA |
| ERBB2 | 0.94786 | 9 | 5 | 4 | 0.3394 | 0 | 0.3422 | 0 | 0.2842 | 0.0005 | NA | NA |
| RELA | 0.94762 | 4 | 8 | 16 | 0.2718 | 0.0006 | 0.2754 | 0.0004 | 0.3455 | 0 | 0.2055 | 0.0065 |
| PPP3R1 | 0.94720 | 9 | 1 | 24 | -0.3697 | 0 | -0.3947 | 0 | -0.3286 | 0.0001 | -0.2931 | 0.0140 |
| FGF9 | 0.94622 | 3 | 2 | 18 | -0.3549 | 0 | -0.3517 | 0 | -0.369 | 0 | NA | NA |
| AKT2 | 0.94588 | 12 | 4 | 7 | 0.2348 | 0.0031 | 0.2613 | 0.0009 | 0.2982 | 0.0003 | NA | NA |
| MEF2C | 0.94579 | 4 | 0 | 24 | -0.3742 | 0 | -0.4007 | 0 | -0.3875 | 0 | NA | NA |

Supplementary Table 6. The 14 genes with cell type specific expression.

| **Symbol** | **Cell_types** |
|---|---|
| GJA1 | astrocytes |
| ERBB2 | astrocytes |
| NOTCH2 | astrocytes |
| FYN | astrocytes |
| EGFR | astrocytes |
| BMPR1A | astrocytes |
| LMNA | astrocytes |
| RELA | astrocytes |
| MEF2C | microglia |
| SNAP25 | neuron |
| FGF9 | neuron |
| KIF3A | neuron |
| HPRT1 | neuron |
| BRCA1 | oligodendrocytes |

Supplementary Table 7. The seven genes with eQTLs located in the transcription factor binding site in the promoter region.

| Ensembl ID | Gene_symbol | SNP | ATAC-seq database | eQTL database | Transcription factor binding motif (in Hocomoco database v11) |
|---|---|---|---|---|---|
| ENSG00000010810 | FYN | rs1057979 | BOCA | Common Mind, GTEx | AP2B_HUMAN.H11MO.0.B, E2F1_HUMAN.H11MO.0.A, KLF9_HUMAN.H11MO.0.C, SP2_HUMAN.H11MO.0.A, RFX1_HUMAN.H11MO.0.B, SP3_HUMAN.H11MO.0.B, KLF3_HUMAN.H11MO.0.B, THAP1_HUMAN.H11MO.0.C, SP4_HUMAN.H11MO.0.A, CTCF_HUMAN.H11MO.0.A, PATZ1_HUMAN.H11MO.0.C, WT1_HUMAN.H11MO.0.C, CTCFL_HUMAN.H11MO.0.A |
| ENSG00000107779 | BMPR1A | rs140745739 | BOCA | GTEx, Common Mind | NR1H4_HUMAN.H11MO.0.B, SP2_HUMAN.H11MO.0.A, SP3_HUMAN.H11MO.0.B, NRF1_HUMAN.H11MO.0.A, ZN341_HUMAN.H11MO.0.C, OSR2_HUMAN.H11MO.0.C, WT1_HUMAN.H11MO.0.C, CTCFL_HUMAN.H11MO.0.A |
| ENSG00000108946 | PRKAR1A | rs8080306 | BOCA | Common Mind | ZN263_HUMAN.H11MO.0.A, MAZ_HUMAN.H11MO.0.A, KLF15_HUMAN.H11MO.0.A, ZSC22_HUMAN.H11MO.0.C, TAF1_HUMAN.H11MO.0.A, ZN467_HUMAN.H11MO.0.C, SP4_HUMAN.H11MO.0.A, ZBT17_HUMAN.H11MO.0.A, VEZF1_HUMAN.H11MO.0.C, ZN341_HUMAN.H11MO.0.C, ZN770_HUMAN.H11MO.0.C, WT1_HUMAN.H11MO.0.C |
| ENSG00000133703 | KRAS | rs7312175 | BOCA | Common Mind | ZN320_HUMAN.H11MO.0.C |
| ENSG00000146648 | EGFR | rs712830 | BOCA | Common Mind, psychEncode | RFX1_HUMAN.H11MO.0.B, TAF1_HUMAN.H11MO.0.A, THAP1_HUMAN.H11MO.0.C, ZN770_HUMAN.H11MO.0.C, WT1_HUMAN.H11MO.0.C |
| ENSG00000160789 | LMNA | rs9427236 | BOCA | Common Mind | MAZ_HUMAN.H11MO.0.A, ZN263_HUMAN.H11MO.0.A, SP2_HUMAN.H11MO.0.A, ZN281_HUMAN.H11MO.0.A, FLI1_HUMAN.H11MO.0.A, ZN467_HUMAN.H11MO.0.C, KLF3_HUMAN.H11MO.0.B, SP3_HUMAN.H11MO.0.B, EGR2_HUMAN.H11MO.0.A, TFDP1_HUMAN.H11MO.0.C, THAP1_HUMAN.H11MO.0.C, ZBT17_HUMAN.H11MO.0.A, VEZF1_HUMAN.H11MO.0.C, WT1_HUMAN.H11MO.0.C |
| ENSG00000221823 | PPP3R1 | rs4519508 | BOCA | Common Mind, psychEncode | MAZ_HUMAN.H11MO.0.A, NR1H4_HUMAN.H11MO.0.B, KLF15_HUMAN.H11MO.0.A, THAP1_HUMAN.H11MO.0.C, PATZ1_HUMAN.H11MO.0.C, SP1_HUMAN.H11MO.0.A |
