## Supplementary Figures for "Genome-wide prediction and integrative functional characterization of Alzheimer’s disease-associated genes"

**List of Supplementary Figures**


| Figures | Description |
|---|---|
| Supplementary Fig. 1 | Comparison between functional enrichment-based method and random sampling method for non-AD (negative) gene selection. |
| Supplementary Fig. 2 | Comparison of the negative controls and randomly selected genes based on their association with AD. |
| Supplementary Fig. 3 | Performances of different brain-region networks based on Random Forest (RF). |
| Supplementary Fig. 4 | Performances of different brain-region networks based on support vector machines (SVM). |
| Supplementary Fig. 5 | Performance of different brain-region networks based on logistic regression (LR). |
| Supplementary Fig. 6 | Validation of the top-ranked genes based on sequence similarity with AD-associated genes. |
| Supplementary Fig. 7 | Validation of the top-ranked genes based on their coexpression with known AD-associated genes. |
| Supplementary Fig. 8 | Validation of the top-ranked genes based on protein-protein interaction networks in the HuRI and STRING database. |
| Supplementary Fig. 9 | Validation of the top-ranked genes based on miRNA-target binding networks. |
| Supplementary Fig. 10 | The correlation of the eigengenes (the first principal component) of Module 3 (a) and Module 4 (b) with three AD traits, namely, CERAD, Braak and CDR scores on the independent MSBB dataset. |
| Supplementary Fig. 11 | The correlation of the eigengenes (the first principal component) of the top-ranked genes with three AD traits. |

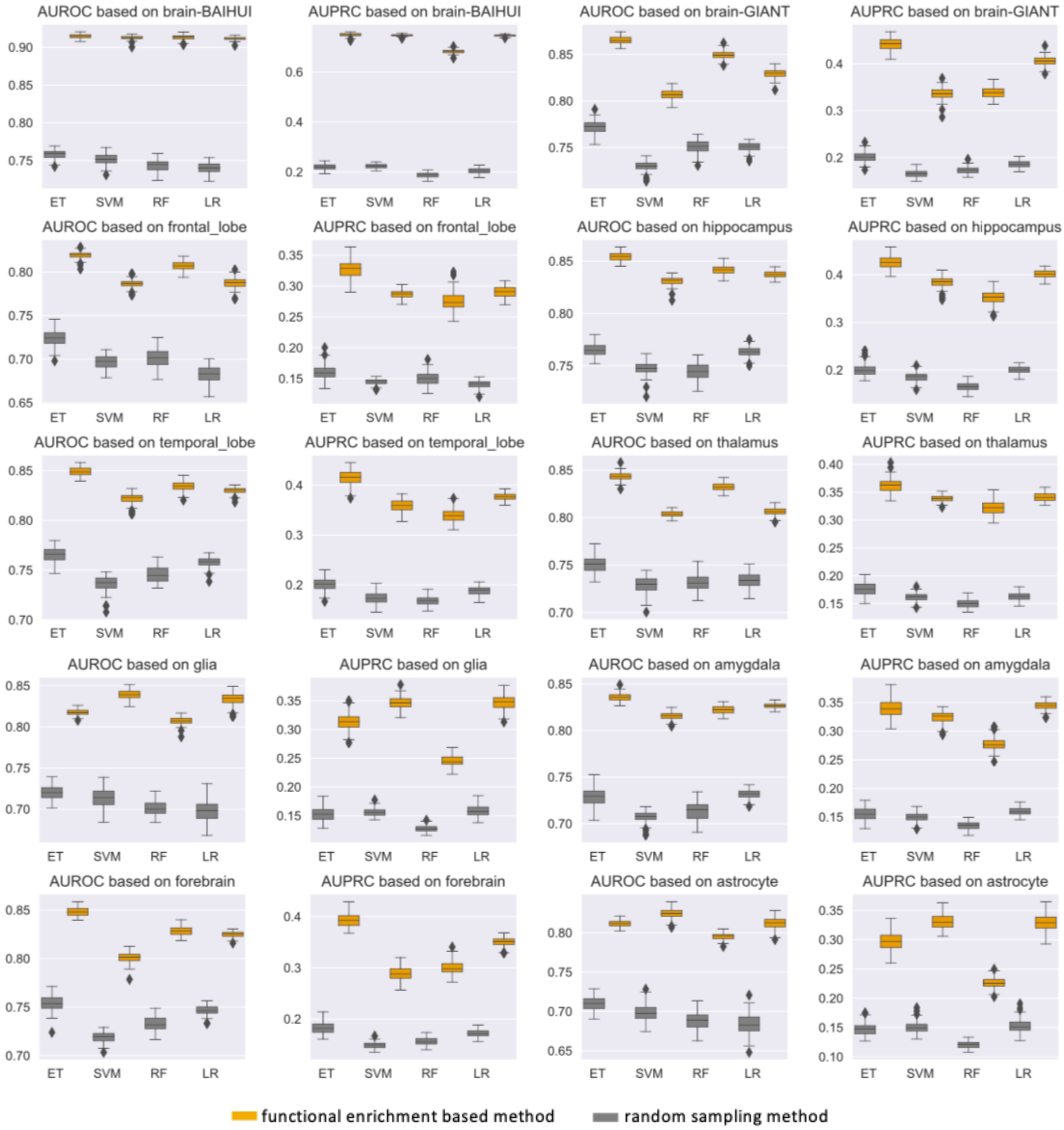


**Supplementary Fig. 1.** Comparison between functional enrichment based method and random sampling method for non-AD (negative) gene selection. The comparison was performed across ExtraTree (ET), random forest (RF), support vector machines (SVM), and logistic regression (LR) models built on the ten brain-specific functional gene networks obtained from GIANT and BAIHUI, respectively.

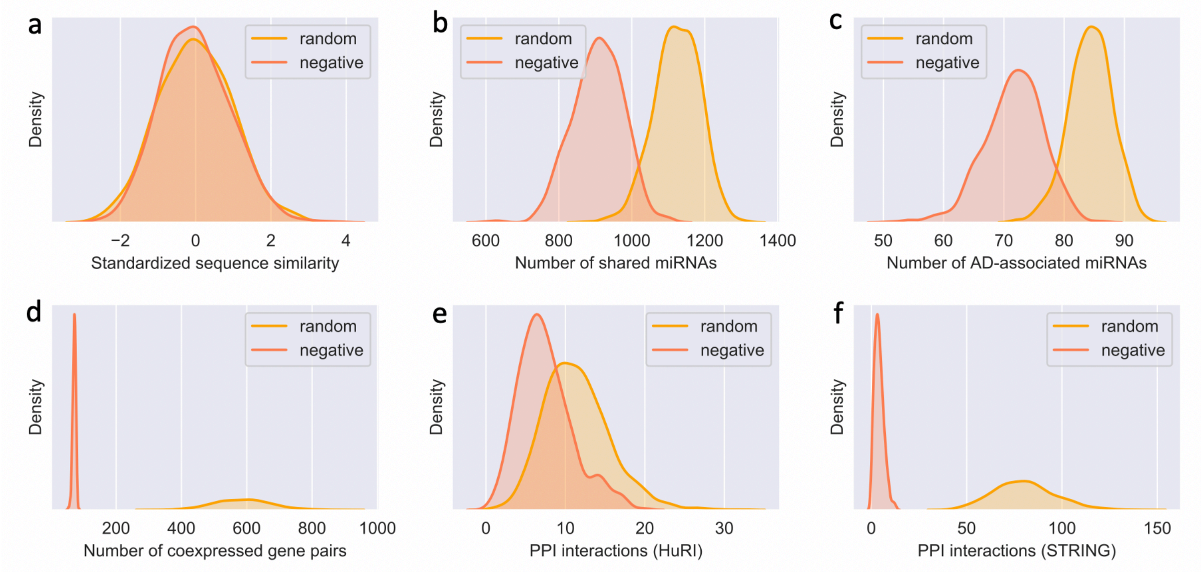


**Supplementary Fig. 2.** Comparison of the negative controls (red) and randomly selected genes (yellow) based on their association with AD. The association is assessed based on the (**a**) Standardized sequence similarity (**b**) Number of shared miRNAs (**c**) Number of AD-associated miRNAs (**d**) Number of coexpressed gene pairs (**e**) PPI interactions (HuRI) (**f**) PPI interactions (STRING).

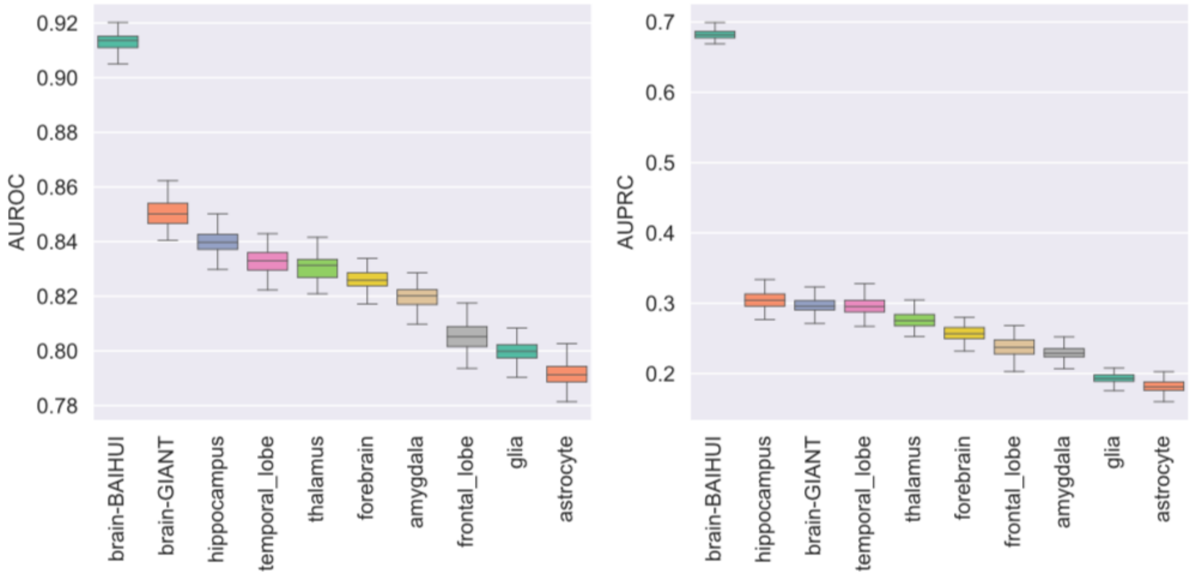


**Supplementary Fig. 3**. Performances of different brain-region networks based on Random Forest (RF).

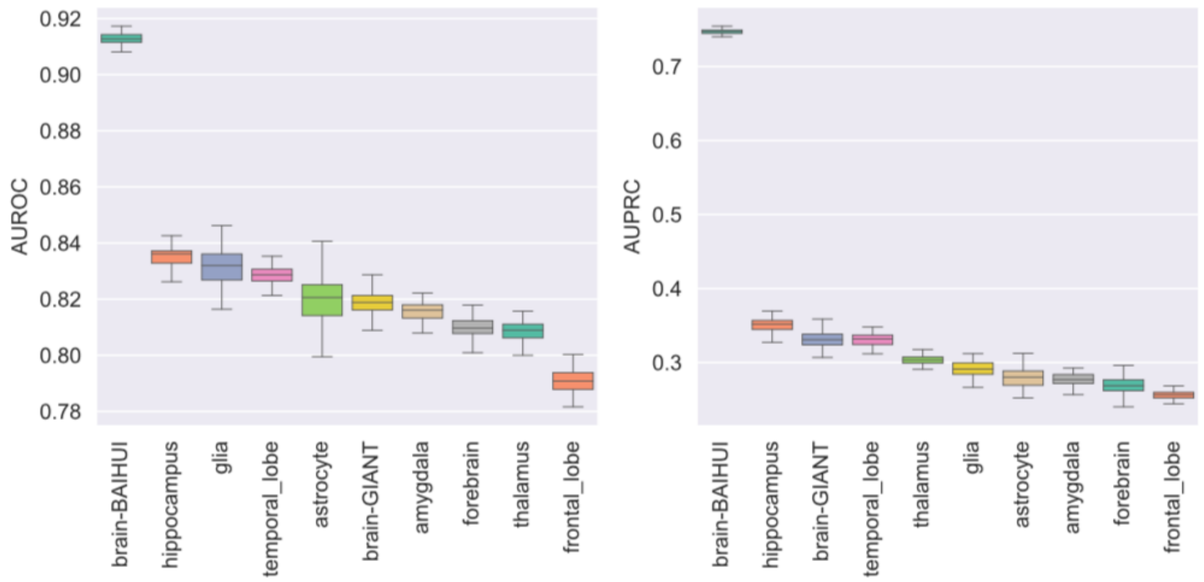


**Supplementary Fig. 4**. Performances of different brain-region networks based on support vector machines (SVM).

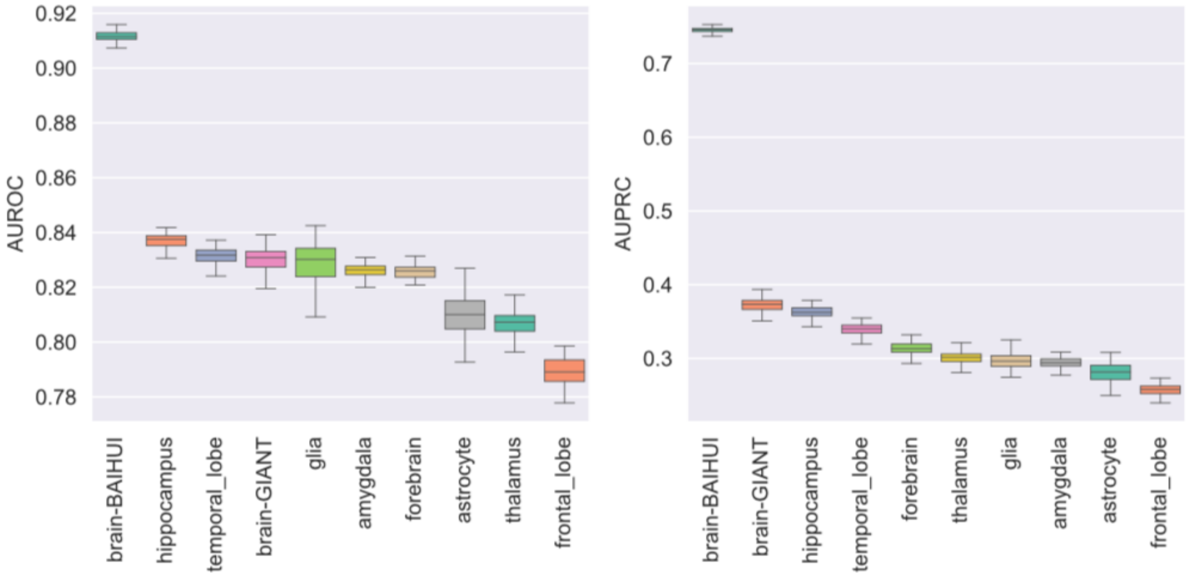


**Supplementary Fig. 5**. Performance of different brain-region networks based on logistic regression (LR).

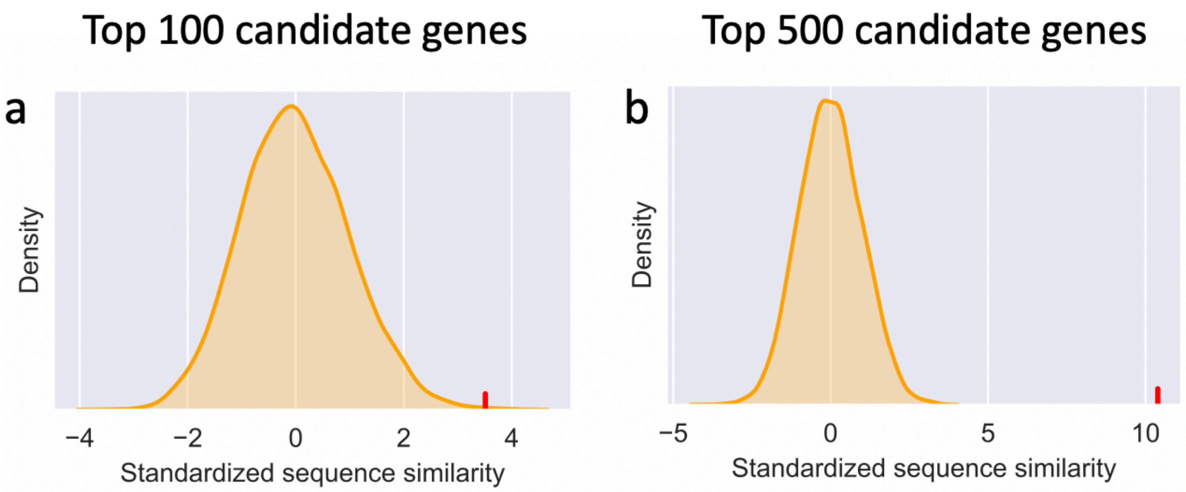


**Supplementary Fig. 6**. Validation of the top-ranked genes based on sequence similarity with AD-associated genes. The average sequence similarities of (**a**) the top-ranked 100 genes and (**b**) the top-ranked 500 genes with AD-associated genes (red vertical line) are significantly higher than that of randomly selected genes (the distribution in yellow). The p-values for (**a**) and (**b**) are 0.0008 and 0.0001, respectively.

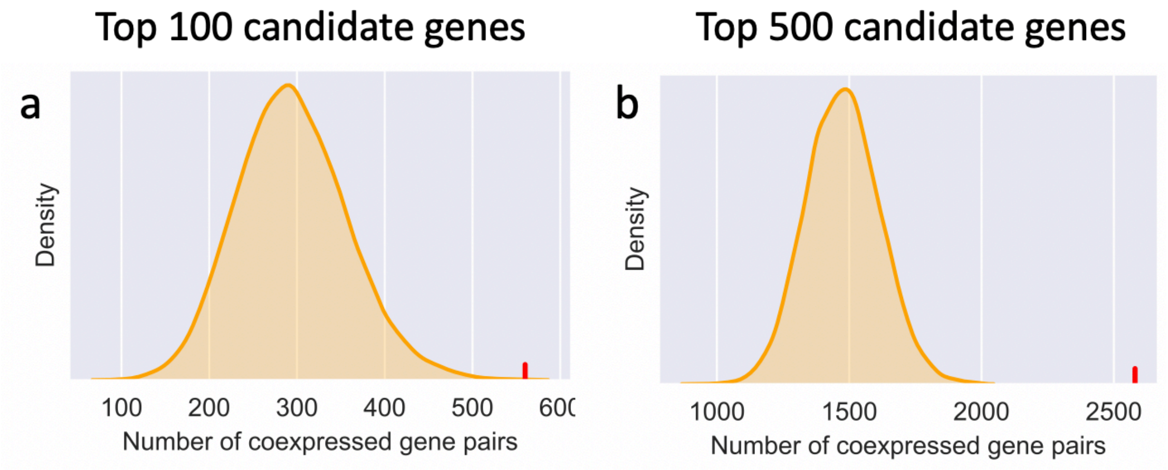


**Supplementary Fig. 7**. Validation of the top-ranked genes based on their coexpression with known AD-associated genes. The number of coexpression gene pairs (Pearson correlation >0.7) of (**a**) the top-ranked 100 genes and (**b**) the top-ranked 500 genes with AD genes (red vertical line) is significantly higher than that of randomly selected genes (distribution in yellow). The p-values for (**a**) and (**b**) are 0.0001 and 0.0001, respectively.

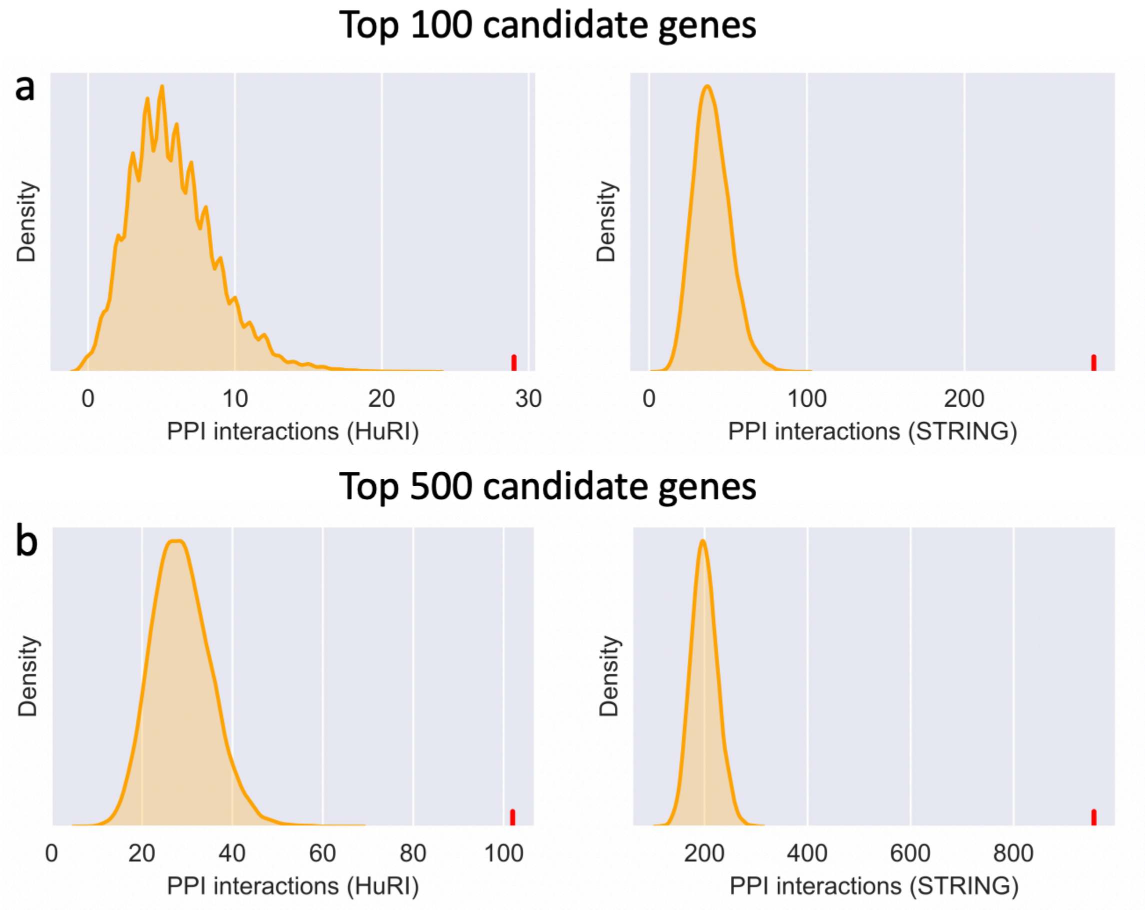


**Supplementary Fig. 8**. Validation of the top-ranked genes based on protein-protein interaction networks in the HuRI and STRING databases. (**a**) The top-ranked 100 genes showed significantly more interactions with the known AD-associated genes (p-value = 0.0001, 0.0001, 0.0001 for the three databases, respectively). (**b**) The top-ranked 500 genes showed significantly more interactions with known AD-associated genes (p-value = 0.0001, 0.0001, 0.0001 for the three PPI databases, respectively). In all plots, the red vertical line and the distribution in yellow denote the result of our predicted genes and randomly selected genes, respectively.

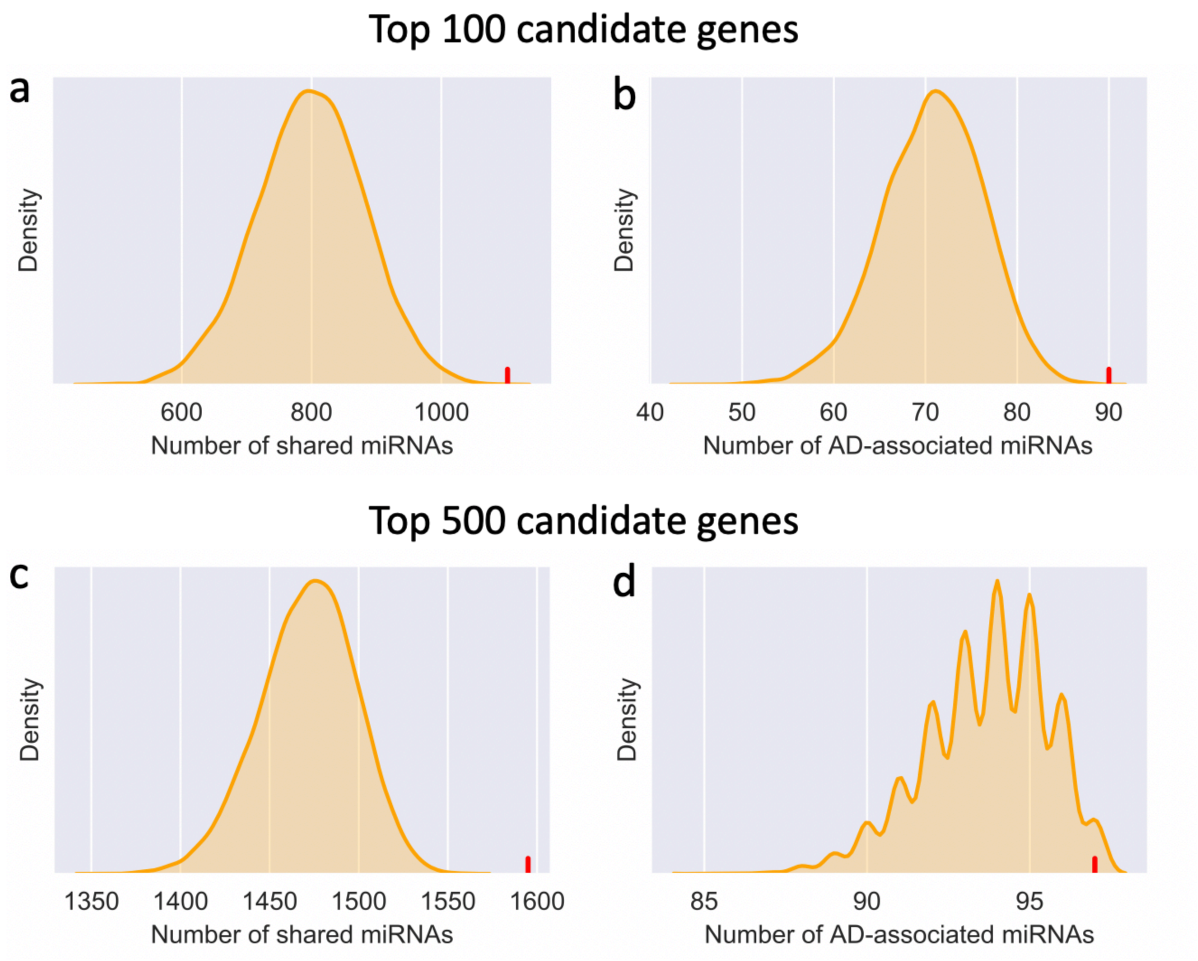


**Supplementary Fig. 9**. Validation of the top-ranked genes based on miRNA-target networks. (**a**) The top-ranked 100 genes share more miRNAs (red vertical line) with AD-associated genes compared to randomly selected genes (yellow distribution) (p-value = 0.0001). (**b**) The top-ranked 100 genes interacted with a significant number of AD-associated miRNAs compared to randomly selected genes (yellow distribution) (p-value = 0.0001) (**c**) The top-ranked 500 genes share more miRNAs (red vertical line) with AD-associated genes compared to randomly selected genes (yellow distribution) (p-value = 0.0001). (**d**) The top-ranked 500 genes interacted with a significant number of AD-associated miRNAs compared to randomly selected genes (yellow distribution) (p-value = 0.0001).

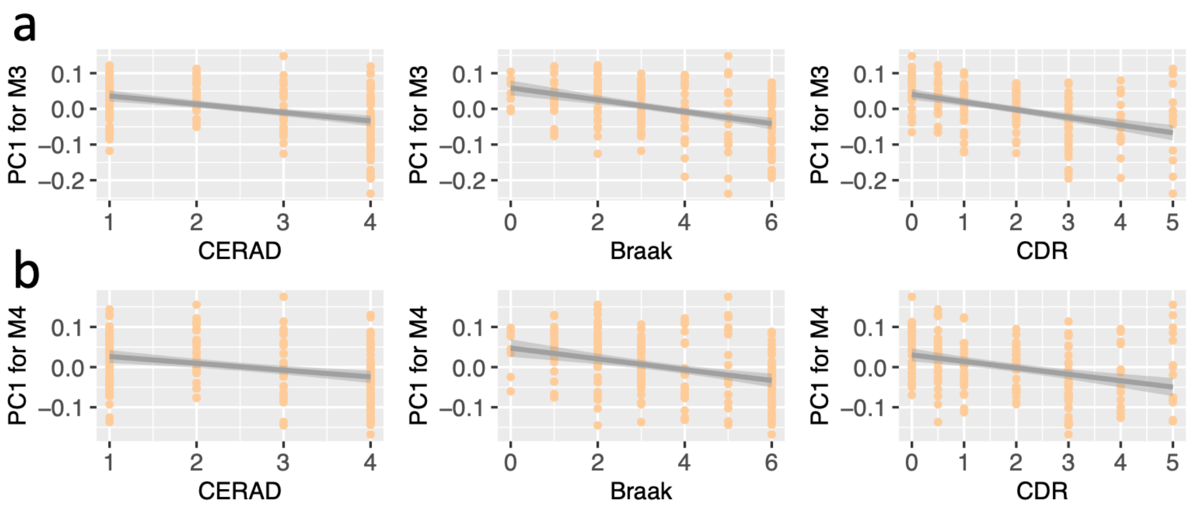


**Supplementary Fig. 10**. The correlation of the eigengenes (the first principal component) of Module 3 (**a**) and Module 4 (**b**) with three AD traits, namely, CERAD, Braak and CDR scores on the independent MSBB dataset.

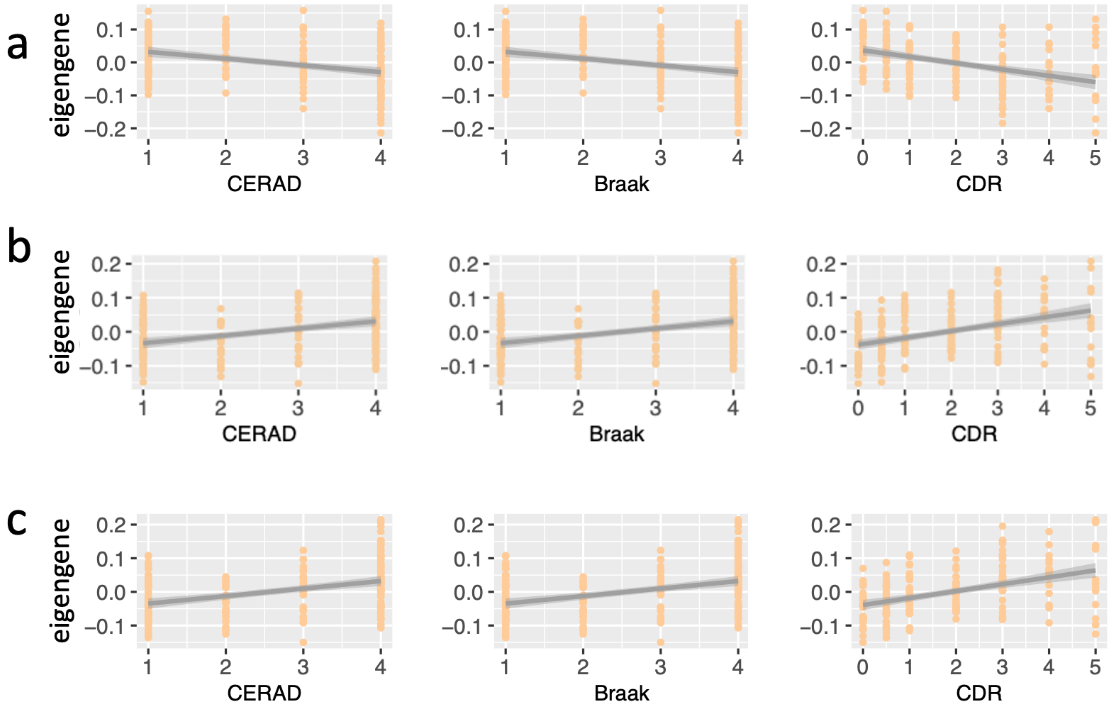


**Supplementary Fig. 11.** The correlation of the eigengenes (the first principal component) of the top-ranked genes with three AD traits. The eigengenes of the top-ranked 100 (**a**), 200 (**b**) and 500 (**c**) genes were significantly correlated with the CERAD, Braak and CDR score (FDR < 0.05 for all the nine correlation tests above). The correlation analysis was performed on the MSBB dataset.
