## Supplementary Note for "Genome-wide prediction and integrative functional characterization of Alzheimer’s disease-associated genes"

### Compilation of AD-associated and non-AD genes

AD-associated (positives) and non-AD (positives) genes are needed to build a machine learning model. First of all, we performed intensive hand-curation to identify confident AD-associated genes (positives) from various disease genes resources, including AlzGene[1], AlzBase[2], OMIM[3], DisGenet[4], DistiLD[5], and UniProt[6], Open Targets[7], GWAS Catalog[8], differentially expressed genes (DEGs) in ROSMAP[9], and published literature. The curated genes from each resource as well as the corresponding criteria were provided in Table 1 in this file. As the AD-associated genes as well as their reliability vary across these resources, we used a vote strategy and selected only the genes that are present in at least two of the above-described resources to ensure higher reliability. In this way, we collected 147 AD-associated genes (Table 2). Of them, most (n=103, 70%) were associated with AD based on GWAS.

Next, we selected a set of non-AD genes, (i.e. negatives), which refer to the genes that have no or minimal association with AD. The main idea of our method for non-AD gene selection was to remove any genes that exhibit potential associations with AD. Let $G_P$ be the set of the 147 collected positives. We then selected a negative gene set $G_N$ containing genes that are potentially associated with AD in the following procedure. We first performed a Gene Ontology (GO) enrichment analysis for the positives using PANTHER[10] and obtained 737 biological process terms (FDR<0.05). It is reasonable to assume that the genes annotated to these GO terms exhibit a potential association with AD compared with a random baseline because they were annotated to the same GO terms (functions) as the positive genes. In addition, we also collected genes that showed any potential association with AD from the above-described resources. These genes were then combined with the genes annotated to the GO terms, forming a set of genes of potential association with AD, denoted by $G_S$. Next, the negative gene set was then calculated as $G_N = G_A - G_S$, where $G_A$ represents all the genes in the network. By removing the 19471 genes identified in $G_S$ from $G_A$ (21122 genes), 1651 genes that were not associated with AD were identified and used as non-AD (negatives) genes.

**Table 1**. Resources for compiling AD-associated genes.

| **Resources** | **Genes** | **Filtering Criteria** | **No. of genes** |
|---|---|---|---|
| OMIM[3] | *PSEN2, HFE, NOS3, PLAU, CALHM1, A2M, PSEN1, ADAM10, MPO, ABCA7, APOE, APP* | All genes associated with AD | 12 |
| DistiLD[5] | *APOC1, BCL3, MS4A4A, FRMD4A, GAB2, MTHFD1L, CR1, EPHA1, CLU, PICALM, SORL1, CD33, ABCA7, CEACAM16, GLIS3, CD2AP, SRRM4, BCHE, ZNF292* | Genome-wide significant association with AD (p-value $< 5.0\times10^{-8}$) | 19 |
| AlzGene[11] | *CLU, TNK1, APOE, SOAT1, PVRL2, TRAK2, CHRNB2, TFAM, CTSD, IL33, THRA, IL1RN, IL10, PICALM, STH, IL6, CD33, TFCP2, VLDLR, SORCS1, IL1B, GAB2, ABCA1, CTNNA3, APOC2, IL1A, BIN1, DAPK1, PON1, CST3, CCR2, GAPDHS, IL8, GRN, ECE1, OLR1, TF, LPL, LRP1, HHEX, GAPDH, LDLR, OTC, ARID5B, TNF, TOMM40, EXOC3L2, PGBD1, PRNP, BCAM, CALHM1, UBQLN1, PCK1, NEDD9, SORL1, ADAM10, MME, ENTPD7, FAS, CH25H, APOC1, ACE, CYP19A1,* | The top ranked AD genes provided on the AlzGene website. | 68 |

| | | | |
|---|---|---|---|
| | *NCAPD2, MTHFR, APOC4, CHAT, CR1* | | |
| AlzBase[2] | *APP, PSEN1, PSEN2, MAPT, APOE, TREM2, PLD3, SORL1, NOTCH3, CLU, PICALM, CR1, ABCA7, BIN1, CD33, CD2AP, MS4A6A, MS4A4E, NME8, EPHA1, PTK2B, SLC24A4, CELF1, FERMT2, ZCWPW1, INPP5D, HLA-DRB1, HLA-DRB5, MEF2C, TP53INP1, CASS4, IGHV1-67* | Top ranked genes supported by genetic evidences | 32 |
| DisGenet[12] | *PSEN1, APP, PSEN2, APOE, SORL1, MAPT, BDNF, IL1B, BACE1, ACE, GSK3B, PLAU* | Genes associated with AD and score > 0.3 | 12 |
| UniProt [6] | *ADAM10, APOE, APP, MT-ND1, MT-ND2, PSEN1, PSEN2, SORL1* | All genes associated with AD | 8 |
| Published literature | *SLC30A6(ZNT6)*[13, 14], *SLC30A4(ZNT4)*[13, 14], *MAOB*[15, 16], *CHRNA7*[17, 18], *IGF1*[19, 20], *VEGFA*[21, 22], *ARC*[23, 24], *PPARG*[25, 26], *BCL2*[27, 28, 29], *IGF2R*[30], *CASP3*[31, 32], *NECTIN2 (PVRL2)*[33, 34, 35], *IGF2*[30, 36, 37], *VSNL1*[38, 39], *ACHE*[40, 41, 42], *CYP46A1*[43, 44, 45], *IDE*[46, 47], *NPY*[48, 49, 50], *IGF1R*[20, 51], *PM20D1*[52], *APBB2*[53, 54], *ESR1*[55, 56, 57], *PAXIP1*[58, 59], *CRH*[60, 61], *SOD2*[62, 63, 64], *SLC2A4*[65, 66], *HMOX1*[67, 68], *DPYSL2*[69, 70], *INSR*[71, 72], *DHCR24*[73, 74, 75], *RELN*[76, 77, 78, 79, 80], *BAX*[81, 82, 83], *REST*[84], *MEOX2*[85, 86], *EIF2AK3*[87, 88], *MAN2A1*[89, 90], *(ABI3, PLCG2)*[91, 92], *DSG2*[26, 93], *MS4A2*[94, 95],*PPP1R37*[96], *RELB*[33, 97], *TREML2*[98], | 1. For GWAS studies: p-value<5.0×10$^{-8}$<br>2. For differential gene expression studies on AD, the gene which is differentially expressed in at least two studies were included. | 56 |

| | | | |
|---|---|---|---|
| | *(HESX1, ADAMTS4, CLNK, HS3ST1, CNTNAP2, ADAM10, APH1B, KAT8, ABI3, ALPK2, ECHDC3, SCIMP, SUZ12P1)*[99] | | |
| GWAS catalog | https://www.ebi.ac.uk/gwas | p-value<$5.0\times10^{-8}$; from both GWAS-reported and GWAS-mapped genes as provided in the GWAS catalog database. | 265 |
| Open Targets[7] | https://www.opentargets.org/ | Genes with association score with AD > 0.5 | 175 |
| ROSMAP | See ref[9] | All DEGs with FDR<0.05 | 2614 |

**Table 2.** The list of 147 AD-associated genes.

| AD-associated genes |
|---|
| *APOE, SORL1, GAB2, CR1, PICALM, CLU, CD33, ABCA7, ADAM10, CD2AP, BIN1, APOC1, TOMM40, INPP5D, PSEN2, EPHA1, APP, MTHFD1L, CNTNAP2, HLA-DRB1, CASS4, BCAM, ABCA1, PTK2B, MS4A6A, FRMD4A, BCL3, SLC24A4, GLIS3, FERMT2, PSEN1, TREM2, ZCWPW1, EXOC3L2, MS4A4A, ACE, APOC4, BZW2, SUCLG2, APOB, SCIMP, SCARB1, RELB, CRY2, PVRL2, CLASRP, ADAMTS4, MMP3, UBE2L3, PPP1R37, ECHDC3, TCF7L2, IL6R, MS4A2, LIPG, MAN2A1, MAPT, ALDH1A2, ABI3, LILRA5, CELF1, PLCG2, HMGCR, OARD1, APH1B, APOC2, OR4S1, STAT4, MS4A4E, PVR, MT-ND2, HS3ST1, CCR2, VASP, CYP8B1, BLOC1S3, PPP1R13L, NFIC, NKPD1, INSR, CNTNAP5, BCAS3, BCHE, BCL2, NME8, CLPTM1, CLNK, UBQLN1, CLMN, IL1B, TRAPPC6A, VSNL1, SORCS1, PPARG, IGSF23, CRH, PSMA1, CHRNB2, FBXL7, CHRNA7, SPON1, MYO16, CHRNA2, VLDLR, KIR3DL2, KIT, HLA-DRB5, BACE1, HLA-DRA, DSG2, CALHM1, RBFOX1, HFE, PILRA, LRP4, HARBI1, TFCP2, CBLC, DPP10, SYNJ1, CDC25B, ACP2, ACHE, PACSIN3, MADD, ZNF652, GSK3B, PFDN1, RIN3, MARK4, GRIN2A, PDGFRB, MAPK8IP1, GRIN3B, CCRL2, ECE1, SCN1A, HBEGF, CACNA1G, CEACAM16, MMP13, ESR1, ALDH5A1, PLAU, SCN8A, CACNA2D1, MMP12* |
